# A Collagen Triple Helix without the Super Helical Twist

**DOI:** 10.1101/2024.09.26.615199

**Authors:** Mark A. B. Kreutzberger, Le Tracy Yu, Maria C. Hancu, Michael D. Purdy, Tomasz Osinski, Peter Kasson, Edward H. Egelman, Jeffrey D. Hartgerink

## Abstract

Collagens are ubiquitous in biology functioning as the backbone of the extracellular matrix, forming the primary structural components of key immune system complexes, and fulfilling numerous other structural roles in a variety of systems. Despite this, there is limited understanding of how triple helices, the basic collagen structural units, pack into collagenous assemblies. Here we use a peptide self-assembly system to design collagenous assemblies based on the C1q collagen-like region. Using cryo-EM we solve a structure of one assembly to 3.5 Å resolution and build an atomic model. From this, we identify a triple helix conformation with no superhelical twist, starkly in contrast to the canonical right-handed triple helix. This non-twisting region allows for unique hydroxyproline stacking between adjacent triple helices and also results in the formation of an exposed cavity with rings of hydrophobic amino acids packed symmetrically. We find no precedent for such an arrangement of collagen triple helices and have designed mutant assemblies to probe key stabilizing amino acid interactions in the complex. The mutations behave as predicted by our atomic model. Our findings, combined with the extremely limited experimental structural data on triple helix packing in the literature, suggest that collagen and collagen-like assemblies may adopt a far more varied conformational landscape than previously appreciated. We hypothesize that this is particularly likely adjacent to the termini of these helices and at discontinuities to the required Xaa-Yaa-Gly repeating primary sequence; a discontinuity found in the majority of this class of proteins and in many collagen-associated diseases.

## Introduction

The collagen family of proteins is a diverse group of macromolecules which form essential supermolecular assemblies in mammals such as collagen fibrils, membrane anchoring fibrils, collagen networks and many others^1^. By definition all collagen and collagen-like proteins have a repetitious Gly-Xaa-Yaa primary sequence that adopt a polyproline type II (PPII) secondary structure^2^. Three either identical (homotrimeric) or non-identical (heterotrimeric) PPII chains further associate into a right-handed superhelix commonly called the triple helix. In the early 1990s the first X-ray crystal structure of a collagen-like triple helix^3^ was solved and since then numerous others have been determined^4,5^. While there have been a number of studies looking at collagen fibril structures using either electron microscopy or X-ray diffraction^6-9^, these studies were all at relatively low resolution and provided limited insight into secondary structure or atomic interactions within the assemblies. Thus, while a number of models have been generated for the structure and packing of a variety of collagenous assemblies, an atomic or near-atomic structural understanding of the interactions that form these structures is missing.

The paucity of structural studies of collagenous assemblies is due to a variety of reasons including the large size of many collagens (thousands of amino acids) as well as the large size and heterogeneity of many collagenous assemblies. While smaller collagenous assemblies exist, many of these suffer from similar issues of heterogeneity as well as difficulty of obtaining enough sample for structural studies. As a result of this, many studies looking at the molecular details of collagenous assemblies have relied upon small chemically synthesized peptides that self-assemble into triple helcies and, in some cases, into higher-order structures. These peptide assemblies have been used for determination of triple helical structures using X-ray crystallography^3,4^ as well as designed assemblies targeted at mimicking the properties of higher-order collagen structures such as fibrils^10-15^.

Another class of collagenous assemblies that these small peptides have been useful for studying are defense collagens^16,17^. These defense collagens are a family of diverse assemblies that all consist of a collagenous structural domain which bind to various immune system targets^18^. One of the of most well-studied defense collagens is complement component 1q (C1q) found in mammalian serum. A key feature of C1q is its “bouquet” structure, containing six flower-like C-terminal globular domains, an intermediate “branching” collagen-like region, and an N-terminal “stem” where the collagen-like portions bundle together (Figure 1a)^19,20^. From previous studies^19-23^ it is known that the collagen-like region in C1q consists of a bundle of six identical triple helices; each triple helix has three distinctive polypeptides, A, B, and C and that the branching region is likely the result of a discontinuity in collagen Gly-X-Y repeat (where X and Y are various amino acids) in chains A and C^21,22,24^.

**Figure 1.**
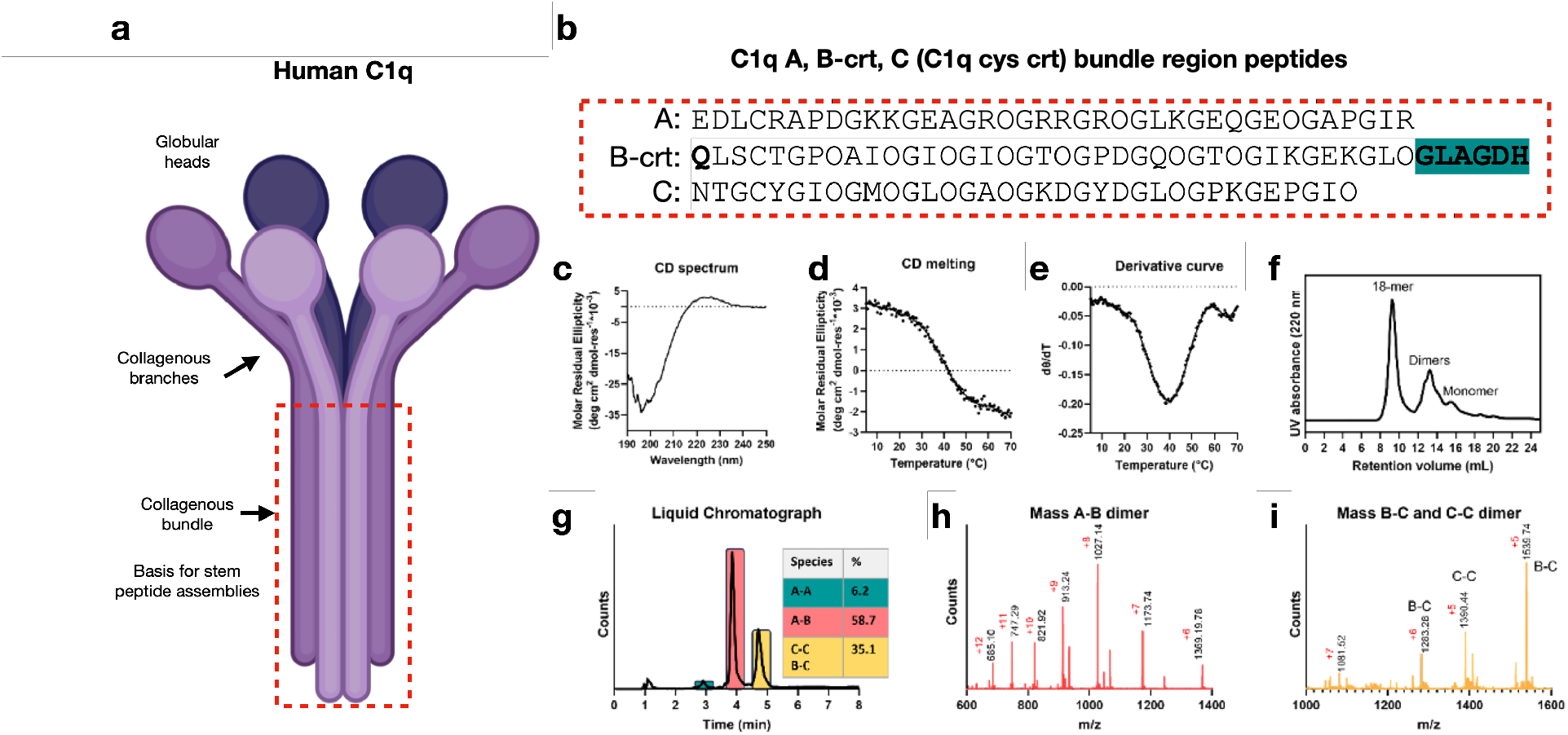
Characterization of peptide assemblies of A, B-Crt and C. **a)** Cartoon depicting the organization of the C1q biological assembly. **b)** Sequence of the three peptides in the C1q cys crt assembly. The terminal Q in peptide B is pyroglutamate. **c)** CD spectrum of the three-peptide mixture. A positive signal at 224 nm is observed. **d)** CD melting curve of the A, B-Crt and C peptide assembly monitored at 224 nm. **e)** The first order derivative of the melting curve displayed in figure giving a melting temperature of 40 °C. d). **f)** SEC of A B-Crt C assembly. **g)** Liquid chromatograph (LC) trace of sample A B-Crt C assembly. **h)** Mass data of the species eluted at 4 mins in the LC trace displayed in g). **i)** Mass data of the species eluted at 4.8 mins in the LC trace displayed in (g).

We have recently recapitulated the unique C1q octadecameric structure in a series of short (∼40 amino acids) synthetic peptides derived from the native sequence of the C1q stem region^16^. Notably, it was observed that the self-assembly of the synthetic system does not necessitate N-terminal disulfide bonds, the C-terminal branching collagen-like region, or the presence of the globular region^16^.The structure of a portion of this assembly was determined to ∼5 Å resolution using cryo-electron microscopy (cryo-EM)^16^. Unfortunately, this low resolution left significant questions about triple helix packing and organization in the assembly unanswered and provided limited insight into the specific stabilizing amino acid interactions in the collagenous assembly.

In this study we synthesized variants of the C1q collagenous stem bundle to optimize cryo-EM imaging. This resulted in a C1q assembly with increased number of particle orientations in cryo-EM grids greatly improving analysis. We determined the structure of a small region of the C1q stem assembly, with a mass less than 30 kDa, to a resolution of 3.5 Å. With this density map we constructed a model delineating the C1q bundled collagen-like region. Our cryo-EM structural analysis of this assembly revealed that the full 70 kDa octadecameric complex had variations in the PPII conformation of the peptide subunits substantially different from those that would be expected for canonical collagen triple helices. In particular, a significant portion of this domain adopts an arrangement of PPII helices lacking the normal superhelical right-handed twist. From this model we delineated pivotal inter-triple helix interactions that foster octadecameric assembly. Additionally, our analysis revealed a configuration wherein hydrophobic amino acid side chains align along the peptide axis, directed towards the lumen. Drawing from this, we synthesized peptide variants featuring varied hydrophobicity and side-chain geometry which substantiated the critical role of the hydrophobic channel in stabilizing the octadecameric assembly.

## Results

### Optimization of the C1q peptide assembly

We initially set out to design peptide assemblies that formed octadecameric C1q stem bundle assemblies that were more suitable for structural determination by cryoEM compared to those previously studied.^16^ All peptide assemblies that were used in this study are detailed in Supplementary Tables S1 and S2. Liquid chromatograms and mass spectra for each of the peptides are shown in Supplementary Figs. S1-12. Eleven peptides were prepared to probe improvements to cryoEM analysis. First, to increase the molecular weight of the assembly we duplicated various portion of the stem sequence in two ways (Peptides A-ext, B-ext, C-ext and A-ext2, B-ext2, C-ext2 as shown in Table S1). While triple helices were observed from these larger assemblies, no significant oligomerization was observed which indicates that rather than stabilizing the oligomer, duplicating these collagen-like sequences actually interferes in some way with the assembly while permitting triple helix formation.

We next attempted to introduce some asymmetry into our structure by slightly increasing the length of peptide B by 6 amino acids at its C-terminus so it could more easily be identified in the density map. This new peptide we call C1q-B “**c**ar**r**ot-**t**op” peptide (B-Crt, as listed in Supplementary Table S1 and Figure 1b). When mixed with the regular A and B peptides, we call their assembly the C1q cys crt assembly (see Figure 1 for assembly scheme, sequence and analysis). Following the mixing of the A, B-Crt, and C peptides (Figure 1b) in buffered conditions containing dithiothreitol (DTT), the assembled peptide structure was characterized using circular dichroism (CD). As illustrated in Figure 1c, the CD spectrum exhibits a characteristic signal indicative of a collagen triple helix at 224 nm. Furthermore, the melting curve of the assembly displays a characteristic thermal transition, with the first-order derivative revealing a melting temperature (*T*_*m*_) of 40 °C (Figure 1d,e). This thermal stability matches our previous study of the native C1q assembly. In our previous study, we established that the formation of this heterotrimeric triple helix is dependent on the composition of the peptide mixture. Specifically, only the tri-peptide mixture folded into a stable triple helix, while binary peptide mixtures or monomeric peptides did not exhibit such stability.^12^

The size exclusion chromatography (SEC) displayed in Figure 1f shows a peak eluting at approximately 9 mL of retention volume, corresponding to the 18-mer assembly. The cluster of peaks eluting at 12-14 mL matches peptide disulfide bonded dimers while the species that elutes at 16 mL matches monomeric peptide. We did not observe significant differences between the new A, B-crt and C sample and the original A, B and C sample in terms of their thermal stability and oligomeric state despite the additional six C-terminal amino acid residues on peptide B.

The A, B-crt, C peptide mixture was equilibrated in the presence of the reducing agent DTT. Nevertheless, liquid chromatography-mass spectrometry LCMS characterization demonstrated that interpeptide disulfide bonds are formed after self-assembly, likely induced by a proximity effect. A-B disulfide bonded dimers were the most prevalent species observed (59%) in addition to significantly smaller quantities of C-C, B-C and A-A dimers (Figure 1g,i and Supplementary Fig. S4.). No Bcrt-Bcrt dimer was observed. This is consistent with previous reports on the disulfide bonding pattern in natural C1q which form A-B and C-C dimers in a final octadecameric (A-B)_6_(C-C)_3_ assembly.^21,23,25^ The presence of small quantities of B-C and A-A dimers in our synthetic system suggest conformational flexibility in the N-terminal region sufficient to permit alternate crosslinking partners.

### Cryo-EM and atomic modeling for C1q cys crt

We collected cryo-electron micrographs of the C1q cys crt assembly (Figure 2a) and upon initial processing of the data were able to produce 2D classes with top views similar to the octadecameric assembly that we showed previously^16^, as well as side views which were not possible previously (Figure 2b). We hypothesize that the six additional residues added to peptide B allowed for better particle behavior in the thin vitreous ice required for optimal cryo-EM imaging. We were first able to determine the structure of the C1q cys crt peptide assembly to low resolution (Figure 2, Supplementary Fig. S13, Supplementary Table S3). This structure was from a subset of particles (∼44,000) and had considerable heterogeneity at the wider ends of the structure revealed by cryoSPARC’s 3D variability analysis (Supplemental Movies S1 and S2). Then we were able to determine the structure of a small narrow region of the assembly to 3.5 Å resolution (Figure 3, Supplementary Figs. S14) from which we were able to build an atomic model of this narrow region (Figure 3, Supplementary Fig. S15). Following this, we were able to use this narrow region model as a starting point to build a model for the full C1q collagenous bundle (Supplementary Fig. S16).

**Figure 2.**
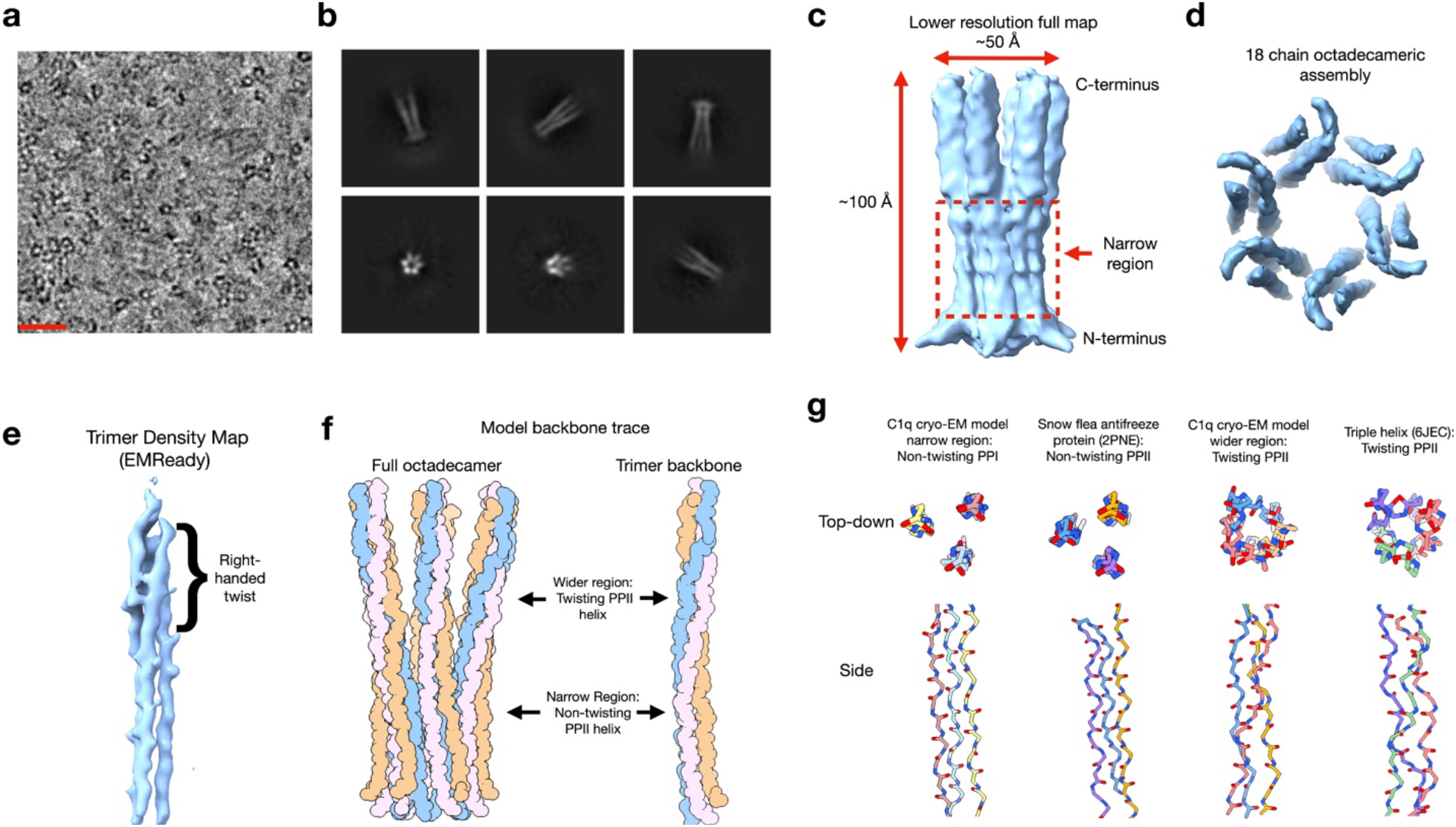
Cryo-EM structure of the full C1q cys crt stem assembly reveals a collagenous assembly with non-canonical variations in PPII triple helical twist. **a)** Cryo-electron micrograph showing the small particles of the stem assembly. The scale bar is ∼200 Å. **b)** Representative 2D class averages of the assembly. **c)** Map of the full C1q stem assembly from the side view showing an oblong structure with a total length of ∼100 Å and a diameter of ∼45 Å. **d)** Map of the full C1q cys crt stem assembly showing a top view where the 18 peptide chains and pore-like nature of the structure is apparent. **e)** Density map of a single C1q stem bundle trimer taken from a version of the full map that has been sharpened using EMready. **f)** Spherical atom representation of a backbone trace into the full C1q cys crt stem assembly density map. The left image shows the full assembly while the right image shows an individual trimer. In the narrow region the subunits in the PPII trimer have no twist with respect to each other while in the wider region the subunits twist around each other in a right-handed fashion typical of collagen triple helices. **g)** Comparison of the non-twisting and right-handed twist PPII conformations of the C1q cys crt assembly with deposited models which are non-twisting or right-handed twisting (triple helix) PPII helices. Note that 2PNE is composed of parallel and antiparallel alignments.

**Figure 3.**
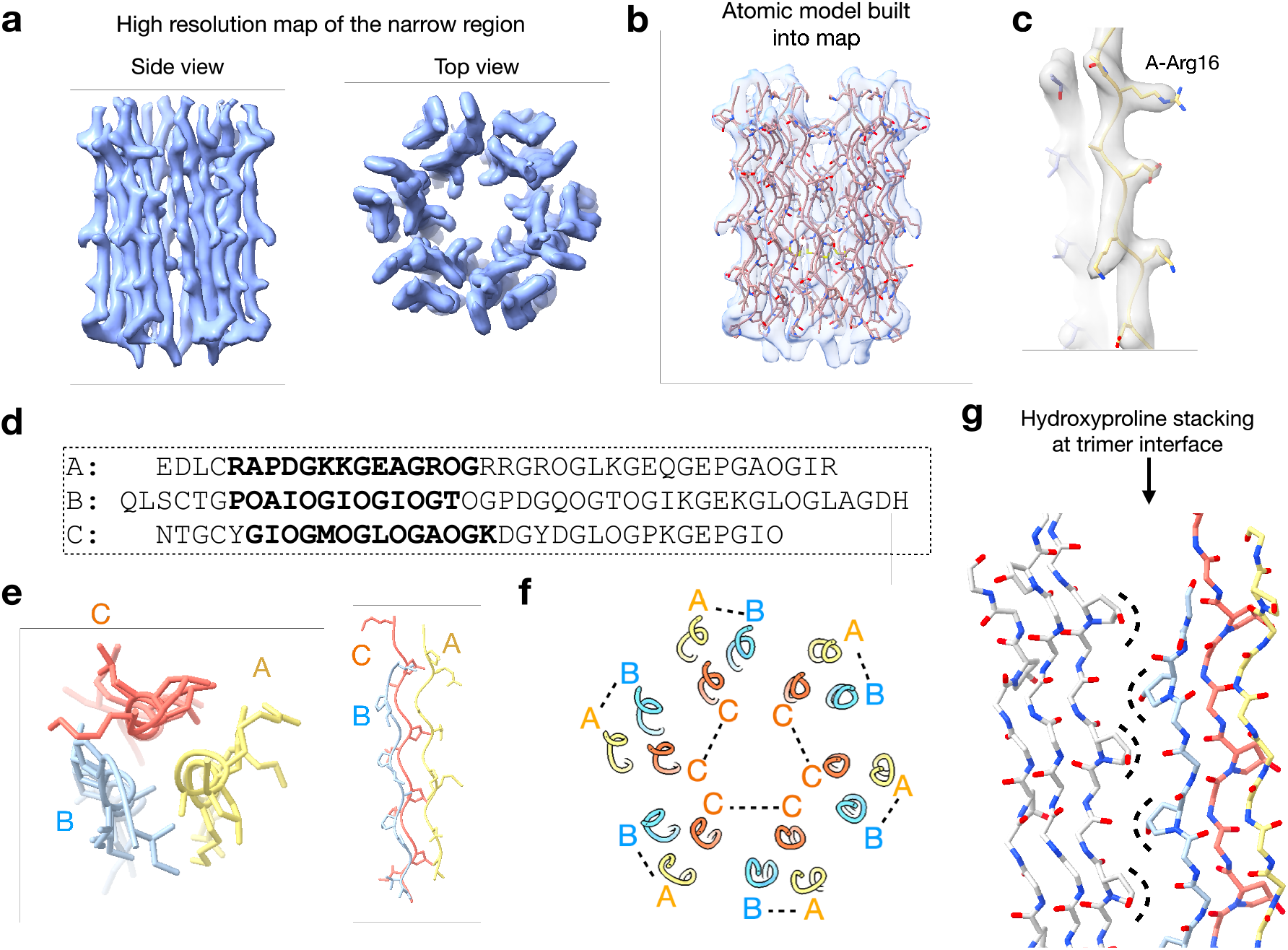
High resolution structure of the narrow region of the C1q cys crt stem assembly reveals the amino acid positioning of the three polypeptide chains. **a)** High resolution density map of the narrow region of the C1q stem assembly from side view (left) and top view perspective (right). **b)** Side view of the narrow region showing an atomic model built into the density map. **c)** Close up on a single trimer showing how the atomic model fits the side chain density. **D**. Full sequence of the three peptides in the C1q cys crt peptide assembly with the sequence corresponding to the narrow, high resolution, region model in bold. **e)** The peptide chains corresponding to peptide A, B-crt, and C are colored yellow, light blue, and orange respectively. The left image shows a top-down view of a single PPII trimer while the right image shows a side view. **f)** The full assembly and where each peptide chain (A,B-crt or C) is positioned in it. The dashed lines indicate likely disulfide bonding. **g)** View of the atomic model for the narrow region of the assembly at the interface between two PPII trimers. Hydroxyproline residues between chain B (light blue) of one trimer interface with hydroxyproline residues from chain C of the adjacent trimer in a fashion resembling the knobs-in-holes packing of alpha-helical coiled coils. The dashed lines are meant to represent the location of each hydroxyproline knob. For simplicity the other amino acid side chains in the model have been hidden.

### C1q collagenous stem structure reveals large variations in triple helix conformation

The low-resolution structure of the C1q cys crt assembly revealed a hollow elongated assembly, ∼100 Å long and ∼30-50 Å wide depending on the region of the structure (Figure 2c). Top-down views of the map revealed the octadecameric nature of the assembly with six copies of the triple-helices (Figure 2d). At the base of the structure there are spokes extending out of the structure, but were at too low a resolution to resolve the individual chains. On the C-terminal side of the spokes the structure is quite narrow and subsequently widens towards the top as shown in Figure 2c. Due to the relatively more hydrophobic nature of the N-terminal sequences, the disulfide bonds, and expected correspondence to the full sized C1q, we hypothesized that the narrow region was the N-terminus. Strikingly, the helical twist of the PPII triple helix changes significantly within this well-defined region. At the narrowest part the three peptide chains in each PPII trimer are nearly parallel to one another and untwisted. As the structure gets wider toward the C-terminus, the peptides wrap around each other with a right-handed twist (Figure 2e) as expected for a canonical collagen triple helix (Figure 2g). Despite the change in superhelical twist, the underlying PPII helicity is preserved throughout. The assembly of three of these PPII helices into a non-twisting, hydrogen bonded system closely resembles the original proposal for the collagen triple helix by Ramachandran in 1954^26^. This non-twisting triple helical region also resembles the PPII packing found in arthropod antifreeze proteins such as that of the snow flea (Figure 2g right two models). The wider region of the C1q stem assembly has a canonical right-handed twist (Figure 2g second from the left model) closely resembling that of existing deposited crystal structures such as that of a portion of the human collagen type II triple helix (Figure 2g far left model).

### AlphaFold3 predictions starkly contrast with experimental structure results

Given the recent release of AlphaFold3^27^ (AF3) and its improved ability to predict both protein secondary structure and protein complexes than its predecessor^28^ we tested to see how reliably it could predict this C1q octadecameric stem bundle assembly. By giving it six copies each of the bundle region of proteins A, B, and C we found that AF3 predicted a model with a hexamer of triple-helical trimers in a long tube-like assembly where the diameter is uniformly about 40-42 Å wide (Supplementary Fig. S17). This is a great improvement over the results that could be obtained with AlphaFold2. But if provided only three copies each of A, B and C, AF3 predicted a trimer of triple-helical trimers, showing that the AF3 predictions in this case are not robust and depend upon the number of chains provided by the user. Further, the register of the three chains in each trimer was wrong as inferred by their inability to form disulfide bonds. In contrast to the AF3 prediction of a uniform diameter, the cryo-EM structure of the C1q cys crt peptide bundle had a narrow N-terminal region and a wide C-terminal region (Figure 2). Also, the AF3 model shows a unform right-handed triple helix conformation for the trimer throughout the structure with exception of the first few apparently disordered N-terminal residues. This contrasts with the presence of the non-twisting N-terminal PPII conformation in the experimental C1q peptide. We tested the performance of AF3 an additional four times by running four independent jobs on the AF3 server (Supplementary Fig. S18). Three out of the four jobs yielded similar results to our first attempt. However, one job failed to predict a hollow octadecameric bundle in four out of the five models it produced (Supplementary Fig. S19).

### Partial structure of human C1q (hC1q) stem bundle validates the peptide assembly system

To validate the biological relevance of our structural findings of the C1q peptide stem assembly we imaged C1q from human serum using cryo-EM (Supplementary Fig. S20). Using SDS PAGE we first confirmed that the C1q components A, B, and C were intact with a molecular weight greater than 25 kDa upon reduction with 2-β-mercaptoethanol, matching the expected molecular weight of a monomer chain. Depending on the gel, the C1q subcomponents can be either a single band^29^ or multiple bands^30^. Under our conditions the three subcomponents were a single band. Therefore, the appearance of the single band for the three C1q subcomponents is expected. We saw 2D-classes that were obvious top-down views as well as apparent side views. However, the side views were much less uniform than those seen for the synthetic C1q cys crt peptide assemblies. For the purposes of this study, we were only interested in the collagenous stem bundle of C1q so our structural results focused on just that aspect. We determined a partial structure of the human C1q stem bundle which displayed similar transitions between a non-twisting PPII helical conformation and a right-handed twisting PPII conformation that were present for the peptide assembly. These results with the human C1q bundle gave us confidence that results with the structure of the C1q cys crt peptide assembly are reflective of the full length natural human C1q stem bundle. Given the presence of both non-twisting and twisting triple helices in the natural C1q cryo-EM structure, we have confidence in our data for the peptide assemblies as reproducing this portion of the natural assembly.

### High resolution cryo-EM structure of the narrow region of the C1q stem assembly

We generated a ∼3.5 Å resolution structure of the narrow region of the assembly (Figure 3a) which resulted in us being able to reliably model the side chains within this region (Figure 3b-c). The sequence of the three chains in the trimer asymmetric unit is shown in Figure 3d and we were able to determine the placement of the three peptide chains in the trimer (Figure 3e) and in the octadecamer as a whole (Figure 3f). From our mass spectrometry data (Figure 1h-i) we observed the canonical A-B and C-C disulfide bonding^22^ in addition to non-canonical C-B disulfide bonding. We show the canonical scheme in Figure 3f, however, the C-B non-canonical bonding would fit the structure we observe as well. The C-C disulfides break the C6 symmetry and the assembly would instead have a lower C3 symmetry. While trying to reconstruct the peptide assembly *ab initio* using C3 symmetry alone (rather than C6 symmetry) was unsuccessful, we did perform a reconstruction with relaxed symmetry (from C6 to C3) using local refinement in cryoSPARC (Supplementary Fig. S21). At the N-terminal base in this structure there was density that could match the C-C disulfide bonding pattern. An interesting structural feature we identified in this model is the interweaving of hydroxyproline residues at the interface of peptide B from one PPII trimer and peptide C from another PPII trimer (Figure 3g) which evokes the “knobs-in-holes” interfaces of α-helical coiled coils^31^. Such an interaction could not exist in a canonical triple helical configuration with a right-handed superhelical twist. The Hyp-Hyp stacking appears to be stabilized by two possible interactions. First, some of the Hyp residues are in optimal positions to hydrogen bond peptide backbone carbonyls and amines. Second, the hydrocarbon portions of the Hyp residues may pack hydrophobically with the hydrocarbons of other Hyp residues as well as hydrophobic residues such as leucine.

It should be noted that the molecular weight of the interpretable portion of the narrow region structure was less than 30 kDa. While the overall molecular weight of the assembly is around 70 Da, we detected a great deal of heterogeneity in our *ab initio* reconstructions of the complex (Supplementary Fig. S14, Supplementary Movies S1 and S2). Since it seems likely that the heterogeneity of the full assembly might hinder the resolution of this more uniform narrow region, it may be reasonable to suggest that this reconstruction is comparable to a cryo-EM structure of a 30 kDa assembly on its own.

### Arginine Mutants Reveal Critical Side Chains for Octadecameric Assembly

We investigated the stabilizing interactions in the octadecamer by making amino acid substitutions in key areas that we identified from our modeling. We noticed that peptide A contains three arginine residues, Arg16, 19 and 22, which exhibit the potential to form hydrogen bonds with the backbone carbonyl oxygen of neighboring triple helices (Figure 4a,f). Examination of our model for the C1q collagenous bundle (Figure 4f) reveals all three of the arginine residues of peptide A are near the interface of PPII trimers (Figure 4g). We can be fairly confident in the side chain positioning of Arg16 as that is well defined in the high-resolution narrow region model (Figure 3c). Given the good backbone fit of the residues into the lower resolution density map (Supplementary. Fig. S16 c,d), we believe that the positioning Arg 19 and Arg 22 are reasonable even though the local resolution in the area of these residues is lower. To determine the importance of this interaction we prepared alanine substitutions of these residues generating new variants designated as A-R16A, A-R19A, A-R22A, and A-tRA, wherein all three arginine residues were substituted with alanine (Table 1). Circular dichroism (CD) of the single substitutions demonstrated assembly into collagen triple helices with comparable thermal stability to the native assembly (Figure 4b-d), however the triple substituted A-tRA assembly displayed significantly decreased thermal stability, with a *T*_*m*_ of 15 °C and SEC showed almost complete elimination of oligomerization (Figure 4e). This suggests the importance of such H bonding networks for the oligomeric stabilization and formation.

**Figure 4.**
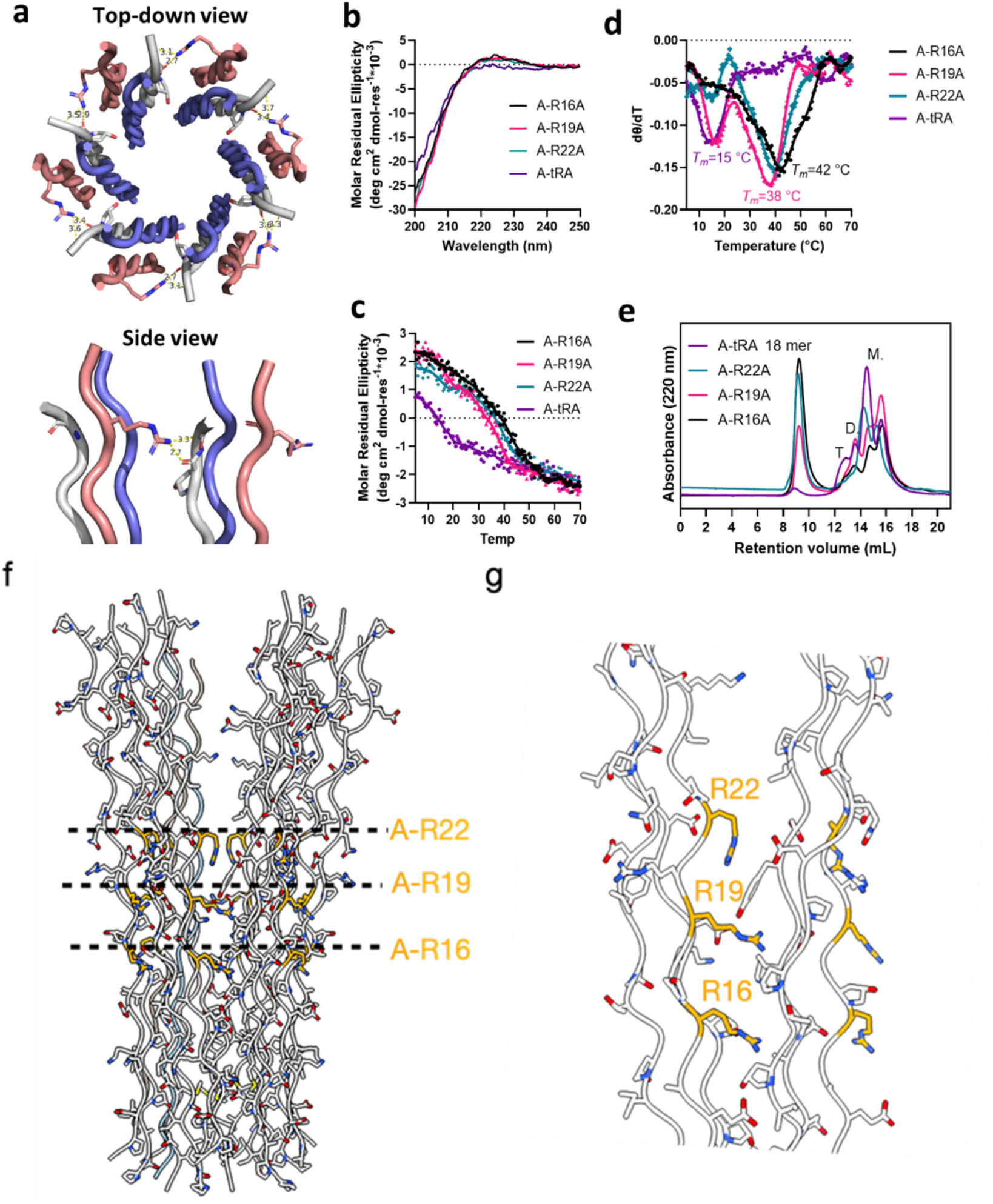
Characterization of C1q cys crt peptide A-arginine to alanine mutant assemblies. **a)** top-down view and side view of the H-bonds between the side chain of the C1q cys crt A-Arg16 residue in one triple helix and the backbone carbonyl oxygen in the other triple helix. For clarity, only two triple helices are shown in the side view. The distance measures between the nitrogen and the oxygen. **b)** CD spectra of the arginine mutants. **c)** CD melting curves of the arginine mutants. Signals were monitored at 224 nm. **d)** First-order derivatives of the melting curves displayed in c). **e)** SEC trace of the arginine mutants. “M.”, “D.” and “T.” represent monomers, dimers and trimers respectively. **f)** Nearly full model of the C1q cys crt complex derived from the high-resolution structure of the narrow region as well as the backbone trace of the wider region. **g)** Close up of the position of A-R16 and approximate positions of A-R19 and A-R22. Since R16 has well defined density in the high-resolution narrow region cryo-EM map we are confident in its placement.

### The C1q Stem assembly contains a cavity with symmetrical hydrophobic rings

Further examination of the model revealed that the lumen in the narrow region was dominated by rings of hydrophobic amino acids (Fig 5a-c). Cross sections of the structure revealed that there were three sets of hydrophobic residues, forming rings with identical residues due to the C6 symmetry at isoleucine 7, methionine 10, and leucine 13 of peptide C (Figure 5b). The diameter across the cavity formed by the hydrophobic rings was ∼5-6 Å (Figure 5c). We found this structural arrangement quite striking because of how the cavity in the narrow region alternated between these hydrophobic rings and hydrophilic spaces between the rings.

**Figure 5.**
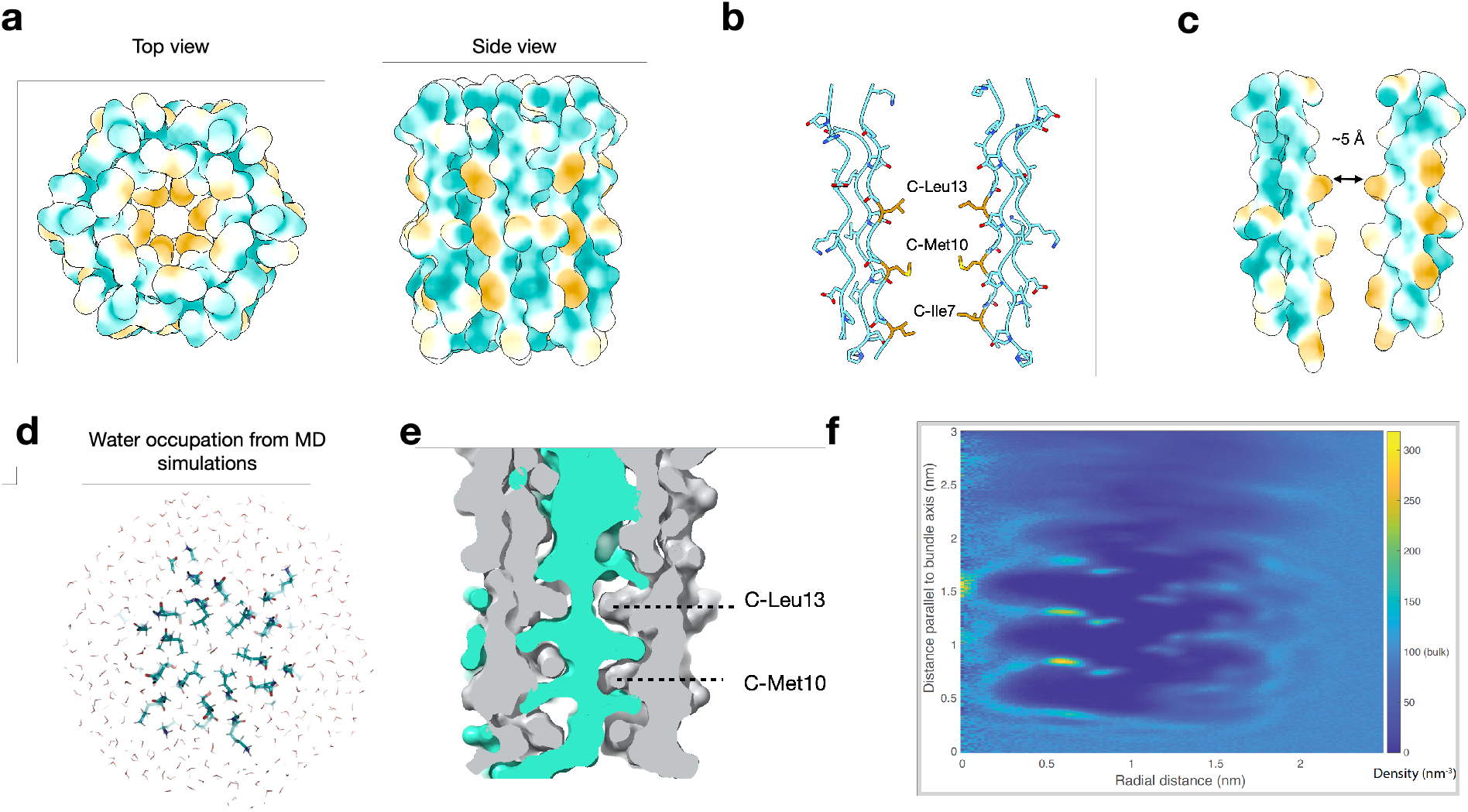
The hydrophobic packing interactions of the C1q stem assembly. **a)** Hydrophobic surface presentations of the narrow region atomic model from a top view. More hydrophobic areas are yellow and more hydrophilic areas are green. Areas in white are neutral compared to the two extremes. **b)** Side view of two opposite PPII trimers highlighting the key chain C hydrophobic residues. **c)** Hydrophobic surface representation of the same region in b. **d**) Results from molecular dynamics simulations showing a top-down view through the hydrophobic region of the pocket showing the lower water occupancy in the central hydrophobic region. **e)** Cross section of the MD simulation results showing both the protein (grey) and water molecules (sea green) as surface representation with van-der Waals atomic radii. The dashed lines point to the approximate positions of the indicated residues. **f)** Graph showing the average solvent density for the larger C1q assembly, showing both solvent depletion in the hydrophobic region and stabilization of specific water sites. Refer to the side bar on the right of the graph for the values of water density that the various colors represent. Very simply, the darkest shades of blue represent areas where the solvent density is less than that of bulk water (less than 100 nm^-3^). Medium blue corresponds to areas of solvent density around that of bulk water (∼100 nm^3^). In increasing order of density areas colored light blue, cyan, shades of green and shades of yellow correspond to areas where the solvent density is greater than that of water

To further our understanding of the hydrophobicity of the narrow region’s cavity, we performed molecular dynamics simulations. Three sets of starting conditions were used, and three replicas were computed for each condition: (1) the narrow region, (2) the larger full atomic model built from the lower resolution structure, and (3) the narrow region with explicit solvent removed from the center of the bundle. Solvent density converged to highly similar results in all three, giving us higher confidence in the results. As shown in Figure 5d-e, the center of the bundle contained water, but the water was highly constricted by both sterics and hydrophobicity (Figure 5e) resulting in as few as 1-2 water molecules in the most constricted regions. Average solvent density was calculated for the full bundle structure averaged over three independent simulations of 500 ns each (Figure 5e) showing reduced solvent density relative to bulk water in the center of the hydrophobic region as well as several “hotspots” of highly ordered water molecules. Similarly, rotational autocorrelation functions were computed for the interior water molecules, and a subset of them had decorrelation times >10x longer than those of bulk water, confirming substantial ordering in the simulation.

Because the MD simulations were performed in the absence of any structural restraints, we can infer that they supported the stability of the narrow region cavity while also yielding insights into the potential behavior of water molecules. We next sought to experimentally test the properties of the hydrophobic region through amino acid substitutions. A well-defined region in the hydrophobic core of the cryo-EM structure corresponded to the density for peptide C methionine 10. We substituted several different kinds of residues into the model at this position (Figure 6a) accounting for a hydrophobic mutation (leucine), a bulky side chain mutation (phenylalanine), a hydrophilic side chain (asparagine), and a charged side chain (aspartate). We made new peptide assemblies with a peptide composition of A, B-crt, and C-M10X with X being one of the aforementioned substitutions. The CD spectra of the peptide assemblies containing C-M10X mutant peptides (Figure 6b) exhibited a characteristic signal at approximately 224 nm, indicative of triple helix formation. We then performed variable temperature CD experiments to characterize the stability of each assembly (Figure 6c-d) and then characterized the mutant assemblies using SEC (Figure 6e). These results show that the assemblies with C-M10L and C-M10N substitutions resulted in octadecameric assemblies (Figure 6e) which were more thermodynamically stable than the “wildtype” peptide assembly (Figure 6c-d). The C-M10F assembly did result in an octadecamer (Figure 6e); however, it showed a broad non-cooperative thermal transition indicating reduced stability and/or heterogeneity (Figure 6c-d). The C-M10D assembly had much reduced thermal stability compared to the other variants (Figure 6c-d) and showed no octadecamer formation (Figure 6e).

**Figure 6.**
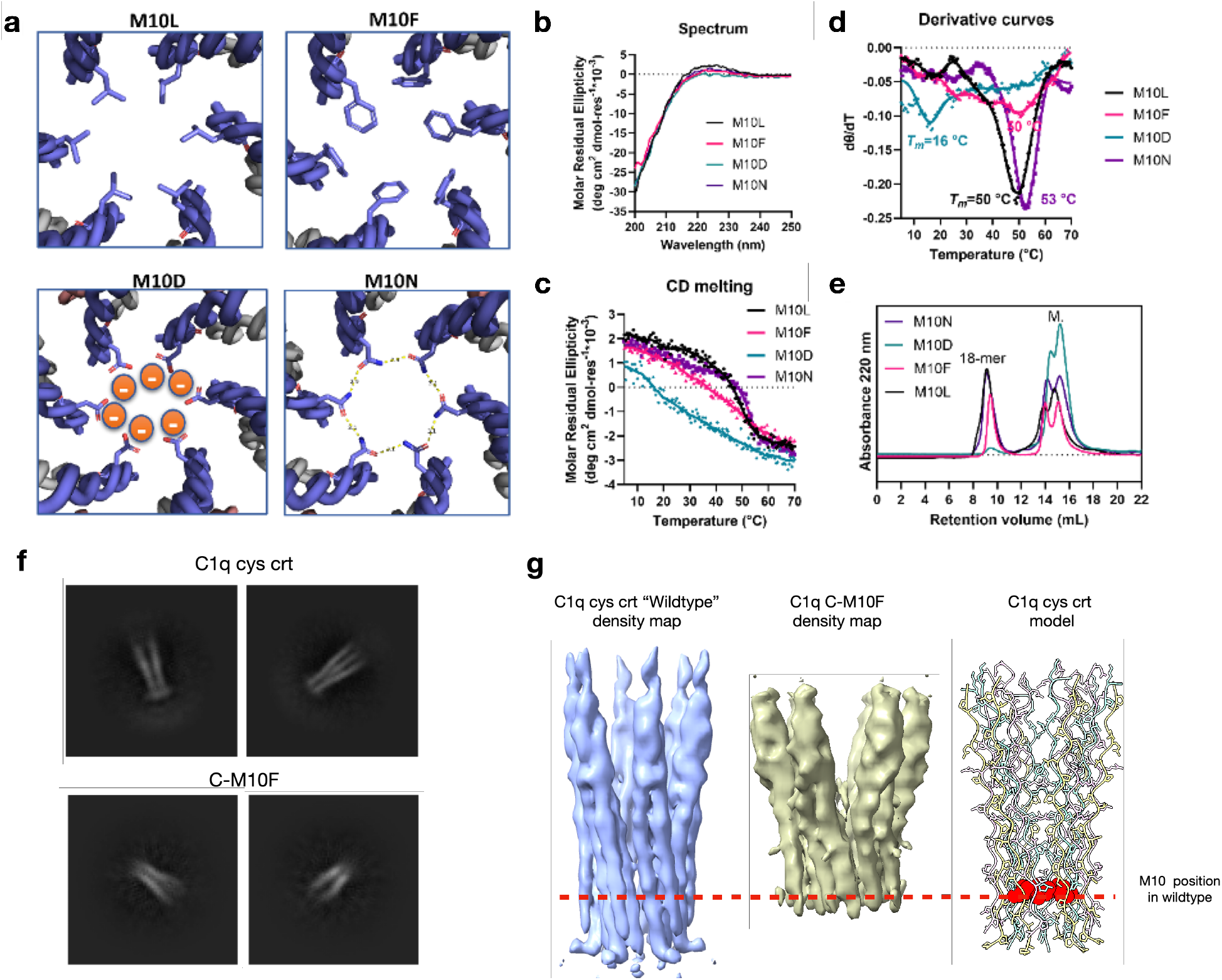
Design and analysis of point mutation variants at the center of the hydrophobic cavity. **a)** Model of the octadecameric assembly of A, B-Crt, and C-M10X. In the model of the octadecameric assembly of M10F mutant, to avoid steric clash, three phenylalanine side chains are pointing toward the C-term and three phenylalanine side chains are pointing toward the N-term in an alternative fashion. In the model of C-M10N assembly, the H bonds measure the distance between nitrogen and oxygen. **b)** CD spectra of A, B-Crt, and C-M10X. **c)** CD melting curves of sample A, B-Crt, and C-M10X, the signal was monitored at 224 nm. **d)** First-order derivative curves of the melting curves displayed in c). **e)** SEC traces of the control assembly of the mutant assemblies. **f)** 2D-Class averages showing side views of the wildtype (top) and C-M10F (bottom) assemblies. **g)** A structural comparison of the full C1q cys crt (“wildtype”) and C-M10F cryo-EM density maps is shown with the C1q cys crt model for reference. The dashed red line represents to position of methionine 10 in the C1q cys crt structure. In the model the methionine 10 side chains are represented by red atoms in spherical representation while the rest of the side chains are shown with atoms in stick representation.

We then used cryo-EM to investigate the structure of the C-M10N assembly and compared it to the “wildtype” C1q cys crt structure (Supplementary Fig. S22). We found that the C-M10N assembly had in general a similar structure to the wildtype one. The only major difference was that the wider C-terminal end of had “spiky” density extending out of the low-resolution region. The C-M10N sample was from a much smaller dataset and the particles underwent less curation than those of the wildtype. Therefore, we suspect that these differences between the two regions are the result of heterogeneity in the M10-N sample. We then turned to solving the C1q C-M10F structure. When compared to the wildtype cys crt assembly most of the side view 2D classes of the C-M10F assembly had a smaller less defined narrow region (Figure 6f). The 3D cryo-EM reconstruction of the C-10F assembly revealed a complex where much of the N-terminal triple helix density was not well resolved when compared to the wildtype C1q cys crt assembly (Figure 6g). We took these structural results to indicate that, as the CD melting and SEC results suggested, the C-M10F does form an octadecameric assembly but its structure is perturbed likely due to steric clashes caused by the phenylalanine substitution.

## Discussion

From a historical stand-point the non-twisting triple helical conformation of the narrow region is an interesting discovery because one of the early models for the triple helix proposed by Ramachandran and Kartha had a strikingly similar packing with three parallel-packed strands adopting a 3_2_ helix^26^. This non-twisting model was abandoned the next year by Ramachandran and Kartha in favor of the three polypeptide chains wrapping around each other adopting a right-handed superhelical twist^32^ and refined that same year in studies by Rich and Crick^33^ and Cowan, McGavin and North^34^. This model for the right-handed triple helix was confirmed with X-ray crystallography nearly 30 years later^3^. However, it appears that an untwisted triple helix has never been shown experimentally until now. Similar untwisted PPII helices are found in PPII bundles which have more than just three chains and adopt a combination of parallel and anti-parallel packing arrangements.^2^ In no case do three such PPII helices arrange in a parallel bundle like we observe here. For a collagenous assembly this packing appears to be a new discovery.

When examining protein structure depositions, we found no examples of PPII or triple helical structures assembling into a hollow structure with “rings” of hydrophobicity. There are many biological structures with hydrophobic pores or pockets for example including both the Orai channel pore as well as thermolysin^35^. Complementing these studies are our molecular dynamics simulations suggesting that the center of the bundle is hydrated but sparsely so in the hydrophobic region. These water molecules in the hydrophobic region also likely have little order and rather than occupying such regions they “pass-through”. Highly ordered water molecules may help stabilize the bundle structure, especially when such molecules engage in hydrogen bonding networks with the peptide backbone in the more hydrophilic regions. Other work has also shown that water closely interacting with a protein surface displays high viscosity and glassy dynamics^36^, which may help act as a “glue” for the bundle.

Overall, we interpreted our C-M10X substitution results as confirmation of our cryo-EM, modeling, and molecular dynamics simulations. This is largely inferred from the fact that the C-M10F structure supported the positioning of our residues in the model while both the C-M10L and C-M10N assemblies formed octadecamer with increased stability. This shows how the pore-like region of our model can support both hydrophobic side chains (Met and Leu) as well as hydrophilic side chains (Asn) as long as they are a certain size. Asn substitutions have been shown to stabilize coiled-coil structures where the satisfaction of hydrogen bonding within the otherwise hydrophobic environment lends specificity to the structure^37^. Additionally, our MD simulations supported the presence of water in constricted fashion in the hydrophobic cavity. The insertion of six Asn residues into the middle of this cavity could increase stability though hydrogen bonding with each other as well as the water molecules. The reduced stability and perturbed structure of the C-M10F assembly support the pore being limited in size. While a C-M10F octadecamer formed, the triple helices before position 10 on peptide C are either very heterogenous in their positioning or not very stable at all. The C-M10F structure also had much less density towards the narrow region where the triple helices were resolvable. This suggests that either the packing interactions or the triple helices themselves before residue 10 in chain C are much less ordered. The C-M10D results are also supportive as the insertion of six negatively charged aspartate residues resulted in the octadecamer assembly being obliterated.

While collagen-based assemblies and their structures have fascinated researchers for over 80 years^38^ few studies have yielded high-resolution structural insights into the packing of triple helices into their supermolecular assemblies. The presence of the non-twisting triple helices observed here suggest that this conformation could be present in other collagen and collagen-like assemblies, none of which currently have atomic structural characterization. We hypothesize that triple helices lacking a super helical twist may be more likely near discontinuities in the (Xaa-Yaa-Gly) as are common in collagens other than types 1-3 as well as near the termini of these repeating units. In the case of the C1q stem bundle the presence of the non-twisting triple helix at the narrow region might be influenced by a variety of factors. First, the non-collagenous N-terminal residues may play a role in triple helix conformation. A lower resolution structure of the partial SP-A peptide assembly appears to have a non-twisting helix for at least a portion of the structure and these peptides also have a non-collagenous N-terminal residues^17^. In this same study it appears that peptides without the non-collagenous N-terminal sequences do not adopt such a non-twisting conformation^17^. However, these structures are quite low resolution in comparison to those presented here and will require future investigation. Another contributing factor to the non-twisting region might be the presence of an alanine in place of a glycine in peptide B position 6. Additionally, the hydroxyproline residues of peptide B and peptide C of interfacing trimers appear to pack into a kind of “knobs into holes” pattern. This Hyp-Hyp stacking ends as the canonical right-handed triple helical structure forms suggesting that this plays some sort of role in stabilizing the non-twisting structure. Lastly, the hydrophobic sequence of many of the N-terminal residues may also play a role. However, it is difficult to discern whether the hydrophobic rings are a cause of the non-twisting triple helical conformation or a symptom of it. One thing that is clear is that without the non-twisting conformation such a hydrophobic packing arrangement is unlikely to occur because with the introduction of a right-handed supercoil the hydrophobic residues of peptide C would no longer all be facing the lumen of the structure. Finally, the lack of super helical twist observed here supports the idea that the stable conformation space available to a collagen triple helix is much larger than previously supposed and this flexibility may be more readily accessible when accounting for packing interactions only present in higher order assemblies.

The difficulty in studying collagenous assemblies is due to a variety of reasons such as heterogeneity, the large size of the supermolecule assemblies, as well as the large size of the collagen molecules themselves^1^. As a result of this, collagen-based peptide assemblies have been quite useful in determining the properties of collagenous assemblies. In the case of the C1q collagenous stem in this paper the main advantages of using synthetic peptides instead of biologically purified or recombinantly produced^30^ assemblies are three-fold. First, we are able to produce peptide assemblies in high quantity and concentration (15 mg/mL) while keeping the critical hydroxyproline residue. Second, we are able to very easily make sequence alterations to probe the properties of the assembly and thus easily test how these alterations affect folding and assembly. Third, we can make slight alterations to the sequences of the peptides of the collagenous stem assembly which only enhance their properties for structural determination. We think that the results in this study highlight the potential for using peptide assembly systems for enhancing the scientific knowledge of collagen structures.

## Methods

### Peptide Synthesis and Purification

All the reagents were obtained from Sigma Aldrich. Peptides were synthesized using solid phase peptide synthesis^39,40^. In detail, a low-loading rink-amide resin was used to generate C-terminal amidated peptides. The resin was swelled twice with dichloromethane (DCM) and twice with N,N-dimethyl formaldehyde (DMF). 25 % v.v. piperidine in DMF was added into the resin to deprotect the FMOC protecting group for 5 mins. The resin was washed five times with DMF and tested with chloranil test or Kaser ninhydrin test to confirm presence of free amines^41^. The amino acid, activating reagent hexafluorophosphate azabenzotriazole tetramethyl uronium (HATU) and a weak base diisopropylethylamine (DiEA) were dissolved in DMF in a molar ration of 1:4:4:6 (resin/amino acid/HATU/DiEA) and mixed for 1 min for pre-activation. The activated amino acid solution was added into the resin for coupling of 20 mins. After coupling, the reaction solution was filtered, and the resin was washed twice with DMF, one time with DCM and one time with DMF.

Negative amine test was observed for each coupling. The deprotection, coupling, washing and testing procedures were repeated for each cycle of amino acid attachment.

After synthesis, the peptides were subject to the final deprotection cycle and washed three times with DCM. The peptides were acetylated with acetic anhydride. In detail, 1:8:4 resin/acetic anhydride/DiEA were mixed and reacted for 45 mins. The acetylation was repeated once. The resin was then washed three times with DCM.

After acetylation, the peptides on the resin were mixed with the cleavage cocktail (90% TFA/2.5% Anisole/2.5% Milli Q H_2_O/2.5% TIPS/2.5% EDT by volume) and reacted for 3 hours. The cleaved peptide solution was drained into a clean round bottom flask. The TFA was blown off with nitrogen gas. Ice-cold diethyl ether was added into the peptide solution to triturate the peptide. The mixture was then centrifuged to obtain the white pellet of the crude peptide. The white pellet was washed once more with cold ether to remove the cleavage cocktail further. After that, the crude peptide was air-dried in the fume hood.

The crude peptides were dissolved in milliQ H_2_O and the solutions were filtered through a 0.2 μm filter before high performance liquid chromatography (HPLC) purification. HPLC was performed in an Empower HPLC instrument equipped with a C18 column (Waters^TM^). The mobile phase A and B are milliQ H_2_O with 0.05% v./v. trifluoracetic acid (TFA) and acetonitrile with 0.05% v./v. TFA, respectively. A general gradient of 10-50% B in 5-32 mins was used for the purification.

### Mass Spectrometry

Mass spectrometry characterization was conducted in an Agilent qToF LC-MS instrument equipped with a diphenyl column. The mobile phase A was milliQ H_2_O with 0.05% v./v. phosphoric acid the mobile phase B was acetonitrile with 0.05% v./v. phosphoric acid. A method with a gradient of 5-75% B from 0-9 mins was used for sample elution.

### Peptide Sample Preparation for Self-assembly

Peptide solutions of 3 mM peptide in 4 mM dithiothreitol (DTT), 20 mM phosphate buffered saline (PBS, pH 7.4) was made for self-assembly. The pH was monitored with a METTLER TOLEDO pH meter (FiveEasy Cond meter F30). The peptide solutions were treated in a 70 °C water bath for 15 mins to denature any pre-formed kinetic species prior to the self-assembly process. The denatured samples were cooled and equilibrated at 4 °C for self-assembly.

### Circular Dichroism

Circular dichroism experiments were conducted in a Jasco J-810 spectropolarimeter equipped with a Peltier temperature controller. Aliquots of the peptides were diluted with 20 mM PBS buffer into a concentration of 0.2 mM for CD measurements. The experiments were performed using cuvettes with a pathlength of 1 mm. The molar residue ellipticity (MRE) value was calculated with the equation, MRE = (θ × m)/(c × l × nr × 10) where θ represents the experimental ellipticity in millidegrees, m is the molecular weight of the peptide (g/mol), c is the peptide concentration (milligrams/milliliter), l is the path length of the cuvette (cm), and nr is the average number of amino acid residues in the peptide. The melting curves were collected with the CD variable temperature measurements, the signals were monitored at the wavelength that gives the maximum MRE value of each sample from 5 °C to 70 °C with a heating rate of 10 °C/hour. The data were processed with Savitzky-Golay smoothing algorithm.

### Size Exclusion Chromatography

SEC experiments were conducted in a Varian Star 3400 instrument equipped with a Superdex Increase 75 30×100 GL SEC column. Aliquots of the peptide solutions were diluted with 20 mM PBS buffer into a concentration of 0.45 mM and 100 μL of 0.45 mM peptide solution was injected for each run. The mobile phase was 20 mM PBS buffer (pH 7.4) and the flowrate was 0.75 mL/min. The signal was monitored at 220 nm with a UV-detector.

### Cryo-EM sample preparation and imaging

All samples for cryo-EM were frozen by applying 3 uL of sample to either a Quantifoil AU 300 mesh grid (1.2 µm/1.3 µm), Quantifoil CU 300 mesh grid (1.2 µm/1.3µm) or Lacey carbon grid (CU 300 mesh). The initial samples were all collected using the quantifoil grids however subsequent collection with the cheaper Lacey carbon grids revealed that particle distributions were either the same as the Quantifoil, if not slightly better. Grids were glow discharged using a GLOQUBE plus^TM^ from Electron Microscopy Sciences applying either a negative voltage or a positive voltage to the grid surface for 20 seconds with a current of 25 mA. Plunge freezing was done using a Vitrobot Mark IV^TM^ with blot forces ranging between 3-8 and blot times between 3-6 seconds. The most optimal condition was a blot force of 5 and a blot time of 5 seconds. All data was collected on a Titan Krios at 300 keV equipped with a K3 direct electron detector using a pixel size of 0.67 Å/pixel and an energy filter of 10 eV. A total dosage of 60 e^-^/Å^2^ was used for each exposure. For the hC1q assembly a high concentration of sample was used making it easy to get plenty of stem bundle structures. Unfortunately, this made identification of the globular heads and collagenous branches difficult.

### CryoEM structural determination of full C1q cys crt assembly as well as M10N, M10F, and human C1q assemblies

For structural determination of the full C1q cys crt assembly (lower resolution) as well as the M10F, M10N, and human C1q samples very similar workflows were used. The workflow for the full low resolution C1q cys crt assembly is shown in Supplementary Fig. S13. After Ab initio reconstruction the best class of structure and corresponding particles were chosen and then a very soft mask was created in cryoSPARC from that structure. The particles, volume, and mask were then used as inputs into homogenous refinement using a maximum alignment resolution of 4-10 Å depending on the structure with static masking. The subsequent outputs from these jobs were then input into a local refinement job with static masking and a maximum alignment resolution from 4-6 Å depending on the job. Additional post-processing steps such as CTF refinement were found to have little effect on the quality of the maps. For presentation of figures in the manuscript noise was removed when present using the map eraser tool in UCSF ChimeraX.

### Cryo-EM Structure of the Narrow region of the C1q cys crt assembly

An overall workflow the cryo-EM structural determination of the narrow region is shown in Supplementary Fig. S14. All image processing and reconstruction steps were performed in cryoSPARC^42^ following a general protocol of motion correction and other processing steps used by us previously^43^ but adapted for smaller particles^16^. Prior to finalizing the workflow for structural determination of the peptide assembly a variety of box sizes were tested resulting in many different structures varying in size. The final box size settled upon was 432 x 432 pixels 2x binned to 216 x 216 pixels. For initial sorting of particles 2D classification was performed where particles with obvious top views and side views were selected and then used as inputs into ab initio refinement. Ab initio refinements using C1, C2, C3, and C6 symmetries were all tested for the octadecameric assemblies with only the C6 symmetry reconstructions yielding reasonable volumes. The failure of the C1 reconstruction to yield an interpretable density map is likely due to the small size (∼70 kDa) and heterogeneity of the data. Using multiple classes for the reconstruction, the various ab initio structures were overall quite heterogenous in size as well as diameter of the wider C-terminal end of the assembly (Supplementary Fig. S14). A single ab initio structural class which appeared to be representative of the overall expected size for the C1q stem bundle assembly and its corresponding particles were chosen for subsequent homogenous refinement with C6 symmetry using a tight mask generated from chosen ab initio volume. Overall, this reconstruction was quite noisy and therefore was lowpass filtered slightly using the gaussian filter function in UCSF ChimeraX^44^ (sDev 1.3). Afterwards, the volume was examined and the wider region of the structure was quite noisy while the narrow region appeared interpretable (Supplementary Fig. S23). A very rough preliminary trace was modeled into the narrow region using coot ^45,46^. From this model a mask was generated in UCSF Chimera^47^ to cut out the good narrow region density. The density of this region was nearly identical to that of the low-pass filtered volume (Supplementary Fig. S25). From this, the narrow region volume was then input into EMReady^48^ for map sharpening. The final EMReady sharpened map was the best overall looking map and the enhanced density appeared to be reasonable given the density present in the non-EMReady-sharpened maps (Supplementary Fig. S23).

### Modeling of the C1q cys crt Narrow Region

The side chain positioning and approximate backbone positioning of the C1q cys crt narrow region closely matched that of the non-twisting PPII snow flea antifreeze protein (PDB: 2PNE)^49^ as well as other non-twisting PPII helcies such as those from the bacteriophage S16 long tail fiber (PDB: 6F45)^50^ as well as the springtail antifreeze protein (PDB: 7JJV)^51^. Based on prior knowledge of the C1q peptide assemblies such as mass spectrometry data, there no evidence for an anti-parallel configuration of the PPII trimers. Thus, three peptide chains from the snow flea antifreeze protein were fit into the C1q cys crt trimer density map individually in such a way to give them a parallel conformation while retaining their hydrogen bonding pattern (Supplementary Fig. S15). Model building was then performed in several steps. First, each side chain density in the narrow region density maps from before and after sharpening by EMReady were analyzed to determine the possible side chains for that corresponding density. Similarly to methods previously published by our lab^52,53^, the absence of large side chain density at a position did not exclude a large side chain, while the presence of large side chain density excluded the possibility of small side chains.

Using the knowledge from our mass spectrometry results showing the disulfide bonding as well as careful threading of each possible sequence permutation through the density maps for each strand of the trimer we found that peptides A and C had one sequence permutation that unambiguously fit into the density map. For chain B there were two sequence possibilities from which we couldn’t completely distinguish from. We chose a possibility consisting of chain B having the following sequence “7-POAIOGIOGIOGT” in the narrow region model over another possibility “4-CTGPOAIOGIOGI” because of a better fit to the raw, low-pass filtered, and EMReady density maps. At this resolution both permutations resulted in a believable scheme for the disulfide bonding between chains A and B and thus the one that matched the EM data was chosen. We took the characterizations of the peptide A arginine mutants (Figure 4) and peptide C methionine mutants (Figure 6) as strong support for our atomic modeling efforts. The models were refined using Phenix Real Space Refinement^54^ and Isolde^55^ for minimization while using coot to keep the Phi and Psi angles as close to the expected values of a PPII helix as possible.

### Model building of the larger C1q Cys collagenous bundle into the low-resolution density maps

To obtain the full model backbones shown in Figure 1 as well as the full model shown in Figure 4, the atomic model of the narrow region obtained from the 3.5 Å resolution map was fit into the lower resolution map of the C1q cys crt assembly. Then the right-handed triple helix peptide 6JEC was fit into the right-handed region of the EMReady full map density map (Supplementary Fig. S16). A single model was made by combining a non-twisting narrow region trimer with the twisting 6JEC trimer fit into their corresponding regions using UCSF Chimera. The sequence of the twisting region was then mutated in coot to match the C1q Cys crt sequence and a full model for the low-resolution map single trimer was generated. 5 symmetrical additional copies were then generated by applying C6 symmetry to the model using the maps coordinate system in UCSF Chimer which resulted in a preliminary structure of the longer sample fit into the full map (Figure 4f-g). In general, the backbone of this model matched the density of both the C1q cys crt (wildtype) and M10N density maps. For the final deposited model a few of the amino acids at the C-terminal end of the structure were deleted because the resolution of the map was too poor to show the backbone and then the model was refined to the map using Isolde^55^ and then Phenix Real Space Refinement^54^.

### Cryo-EM density map resolution assessment

The map:map Fourier shell correlation curves are shown in Supplementary Fig. S24. The map:model FSC curves for the structures where a model was built into are shown in Supplementary Fig. S25. At first glance, the 3.5 Å resolution estimate at 0.5 map:model FSC generated by the narrow region model and the EMReady sharpened density map is surprising. However, we then took the narrow region density map (not sharpened by EMReady), generated a mask with it in cryoSPARC, and subsequently used that to generate the map:map FSC of the volume used for the narrow region structure (Supplementary Fig. S24a yellow map). The resulting 0.143 “gold-standard” map:map FSC estimate was 3.5 Å resolution.

### SDS-PAGE analysis of human serum C1q assemblies

Human serum C1q was purchased from Innovative^TM^ Research. The standard procedure for SDS-PAGE was followed using Mini-PROTEAN precast gels with a 4-20% gradient and tris/glycine/SDS buffer. The hC1q sample was mixed at a 1:2 ratio with 2x Laemeli Sample Buffer from Bio-Rad. For the reducing conditions the Laemeli sample buffer also contained 2-β-mercaptoethanol. After addition of the lamelli sample buffer, the samples were boiled for 5-10 min at 100°C and then loaded onto the gel. The gels were also loaded with Precision Plus Protein Dual Color Standards. The gels were run at 130 V until the protein standard lanes were resolved to satisfaction and then stained with Coomassie Brilliant Blue R-250 staining solution from Bio-Rad. Gels were de-stained with the Coomassie Brilliant Blue R-250 de-staining solution.

### Molecular dynamics simulations

Structures were prepared for molecular dynamics simulation using the CHARMM-GUI interface^56^. Simulations were performed using GROMACS 2023^57^ with the CHARMM36^58^ parameter set, explicit TIP3P water, and 150 mM NaCl. Simulations used Particle Mesh Ewald long-range electrostatics, a velocity-rescaling thermostat^59^ at 37 ºC, and Parrinello-Rahman pressure coupling^60^ at 1 bar. Three replicas were run of each simulation condition of length ≥500 ns each. Convergence of water density plots across the different starting conditions was used as an indicator of adequate sampling.

## Supporting information

Supplemental Experimental Data

## Acknowledgments

This work was supported in part by the NSF CHE (grant number 2203937), The Robert A. Welch Foundation (Grant C-2141), and the NIH NIGMS (grant numbers GM122510 and GM138444). Computational resources were provided by the National Academic Infrastructure for Supercomputing in Sweden (NAISS). Thanks to Arghadip Dey for assistance with peptide synthesis.

