## Supplemental Experimental Data for "A Collagen Triple Helix without the Super Helical Twist"

[1] Department of Biochemistry and Molecular Genetics, University of Virginia School of Medicine, Charlottesville, VA, 22903, USA
[2] Department of Chemistry, Rice University, 6100 Main Street, Houston, TX, 77005, USA
[3] Molecular Electron Microscopy Core, University of Virginia School of Medicine, Charlottesville, VA, 22903, USA
[4]Center for Advanced Research Computing, University of Southern California, Los Angeles, CA 90089
[5]Departments of Chemistry & Biochemistry and Biomedical Engineering, Georgia Institute of Technology, Atlanta, GA 30332
[6] Department of Bioengineering, Rice University, 6100 Main Street, Houston, TX, 77005, USA
[†] These two authors contributed equally to this work, listed alphabetically.

### Supplementary Information

***Table S1. Sequences of the Collagen-like Peptide Investigated in this Study.***

| Peptide | Sequences |
|---|---|
| A | EDLCRAPDGKKGEAGROGRRGROGLKGEQGEOGAOGIR |
| B | **Q**LSCTGPOAIOGIOGIOGTOGPDGQOGTOGIKGEKGLO |
| C | NTGCYGIOGMOGLOGAOGKDGYDGLOGPKGEPGIO |
| A-ext. | EDLCRAPDG**KKGEAGROGRRGROGKKGEAGROGRRGROG**LKGEQGEOGAOGIR |
| B-ext. | **Q**LSCTGPOA**IOGIOGIOGTOGPDGIOGIOGIOGTOGPDG**QOGTOGIKGEKGLO |
| C-ext. | NTGCYG**IOGMOGLOGAOGKDGIOGMOGLOGAOGKDG**YDGLOGPKGEPGIO |
| A-ext.2 | EDLCRAPDGKKGEAGROGRRGRO**GLKGEQGEOGLKGEQGEO**GAOGIR |
| B-ext.2 | **Q**LSCTGPOAIOGIOGIOGTOGPD**GQOGTOGIKGQOGTOGIK**GEKGLOGLAGDH |
| C-ext.2 | NTGCYGIOGMOGLOGAOGKDGYD**GLOGPKGEPGLOGPKGEP**GIO |
| B-crt | **Q**LSCTGPOAIOGIOGIOGTOGPDGQOGTOGIKGEKGLO**GLAGDH** |
| A-R16A | EDLCRAPDGKKGEAG**A**OGRRGROGLKGEQGEOGAOGIR |
| A-R19A | EDLCRAPDGKKGEAGROG**A**RGROGLKGEQGEOGAOGIR |
| A-R22A | EDLCRAPDGKKGEAGROGRRG**A**OGLKGEQGEOGAOGIR |
| A-tRA | EDLCRAPDGKKGEAG**A**OG**A**RG**A**OGLKGEQGEOGAOGIR |
| C-M10D | NTGCYGIOG**D**OGLOGAOGKDGYDGLOGPKGEPGIO |
| C-M10N | NTGCYGIOG**N**OGLOGAOGKDGYDGLOGPKGEPGIO |
| C-M10L | NTGCYGIOG**L**OGLOGAOGKDGYDGLOGPKGEPGIO |
| C-M10F | NTGCYGIOG**F**OGLOGAOGKDGYDGLOGPKGEPGIO |

*"ext." represents "extended". "O" represents 2S,4R-hydroxyproline. The glutamine "Q" residue at the N-terminus of peptide C1qB and its variant peptides is 5-oxopyrrolidine-2-carboxylic acid. The colored region indicates mutation positions compared to the A, B, C peptides that modeled the native C1q. Peptides A, B, and C were designed and investigated in the reference.*[12]

***Table S2 Peptide sequence and molecular weight information.***

| Peptide | Sequences | $[M+H]^+$ expected | $[M+H]^+$ observed |
|---|---|---|---|
| A | EDLCRAPDGKKGEAGROGRRGROGLKGEQGEOGAOGIR | 3990.1 | 3991.7 |
| B-crt | **Q**LSCTGPOAIOGIOGIOGTOGPDGQOGTOGIKGEKGLO**GLAGDH** | 4222.0 | 4221.1 |
| C | NTGCYGIOGMOGLOGAOGKDGYDGLOGPKGEPGIO | 3474.6 | 3476.5 |
| A-R16A | EDLCRAPDGKKGEAG**A**OGRRGROGLKGEQGEOGAOGIR | 3944.0 | 3946.0 |
| A-R19A | EDLCRAPDGKKGEAGROG**A**RGROGLKGEQGEOGAOGIR | 3944.0 | 3946.0 |
| A-R22A | EDLCRAPDGKKGEAGROGRRG**A**OGLKGEQGEOGAOGIR | 3944.0 | 3945.9 |
| A-tRA | EDLCRAPDGKKGEAG**A**OG**A**RG**A**OGLKGEQGEOGAOGIR | 3773.9 | 3775.9 |
| C-M10D | NTGCYGIOG**D**OGLOGAOGKDGYDGLOGPKGEPGIO | 3456.6 | 3458.6 |
| C-M10N | NTGCYGIOG**N**OGLOGAOGKDGYDGLOGPKGEPGIO | 3455.6 | 3456.6 |
| C-M10L | NTGCYGIOG**L**OGLOGAOGKDGYDGLOGPKGEPGIO | 3454.6 | 3456.6 |
| C-M10F | NTGCYGIOG**F**OGLOGAOGKDGYDGLOGPKGEPGIO | 3488.6 | 3490.6 |

*"ext." represents "extended". "O" represents 2S,4R-hydroxyproline. The glutamine "Q" residue at the N-terminus of peptide C1qB and its variant peptides is 5-oxopyrrolidine-2-carboxylic acid.*

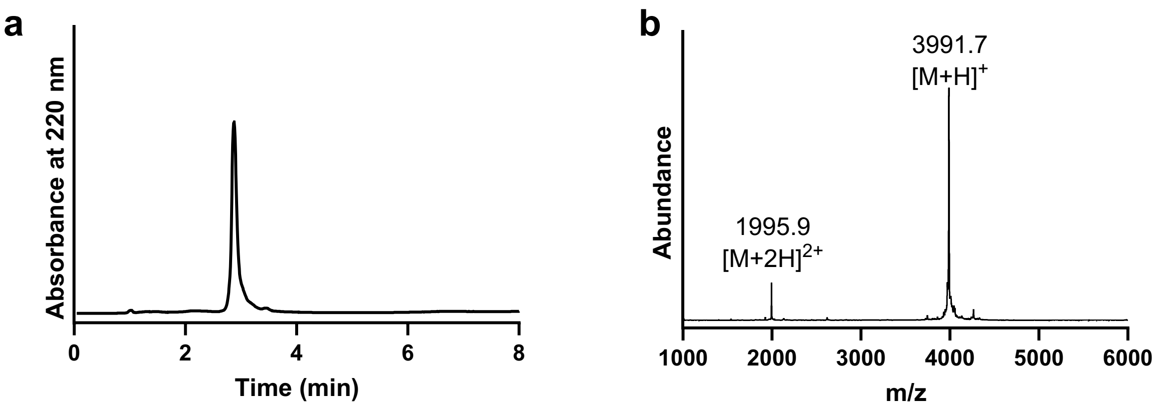


**Supplementary Figure S1.** a) liquid chromatography of pure peptide A. b) mass spectrum of pure peptide A.

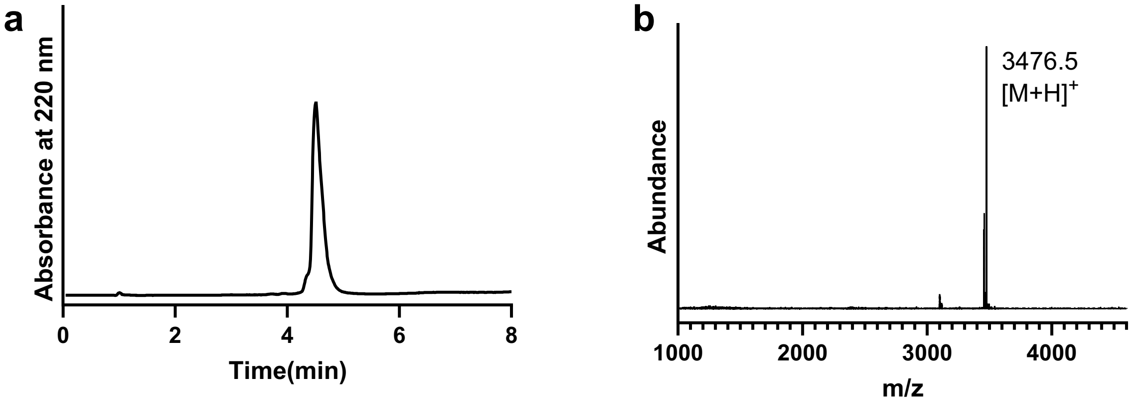


**Supplementary Figure S2.** a) liquid chromatography of pure peptide C. b) mass spectrum of pure peptide C.

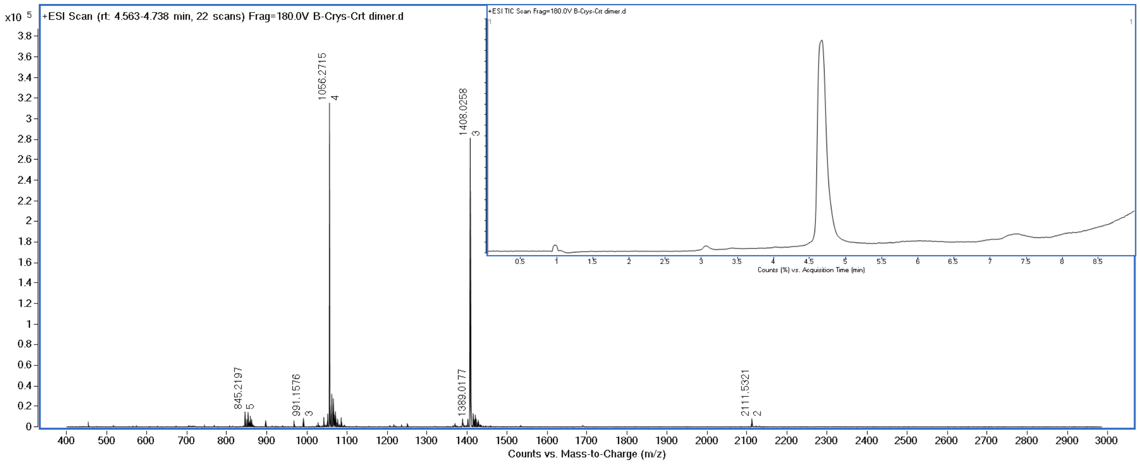


**Supplementary Figure S3.** LC-MS characterization of C1qB-Crt peptide.

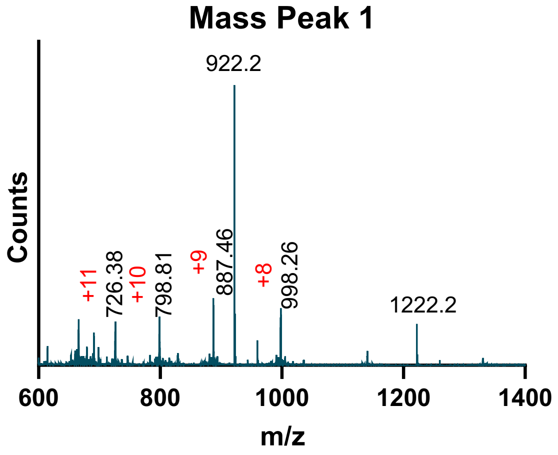


**Supplementary Figure S4.** Mass spectrometry of the first peak in the liquid chromatography displayed in Figure 1 e. The molecular weight observed corresponds to the A-A dimer. The peaks of 922.2 and 1222.2 Da are the calibration peaks of protein standards.

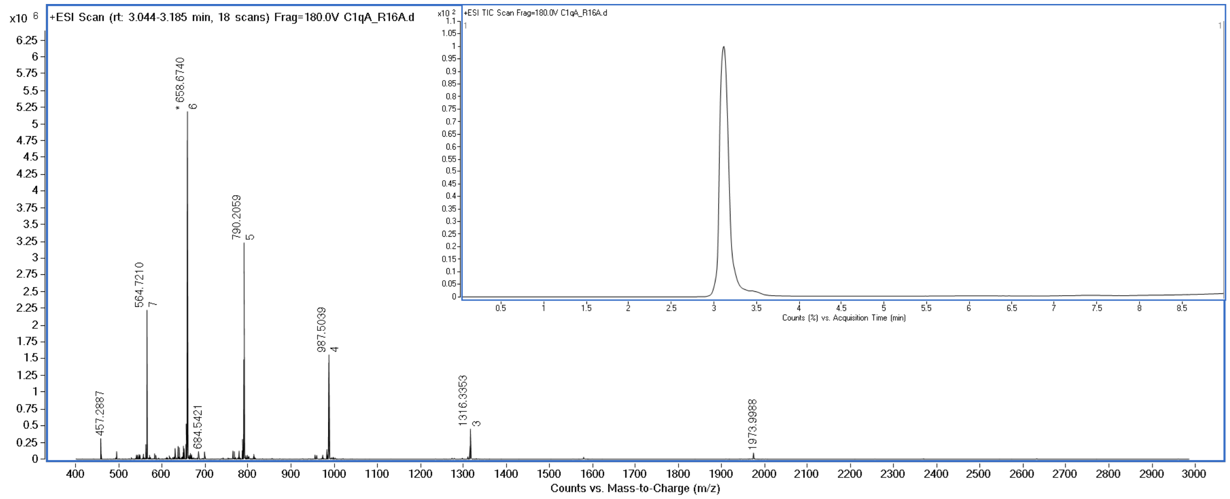


**Supplementary Figure S5.** LC-MS of C1qA-R16A peptide.

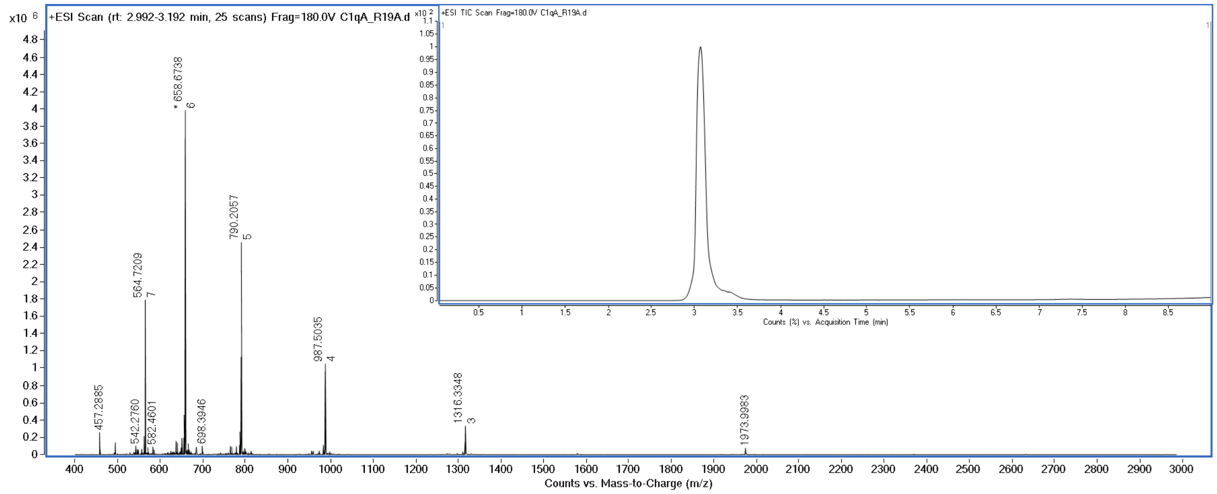


**Supplementary Figure S6.** LC-MS of C1qA-R19A peptide.

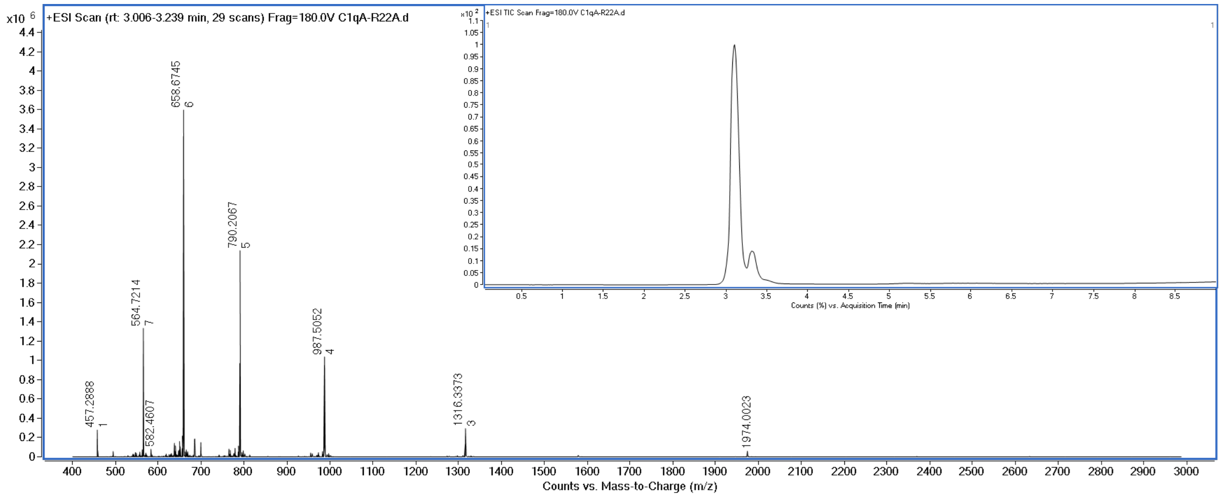


**Supplementary Figure S7.** LC-MS of C1qA-R22A peptide.

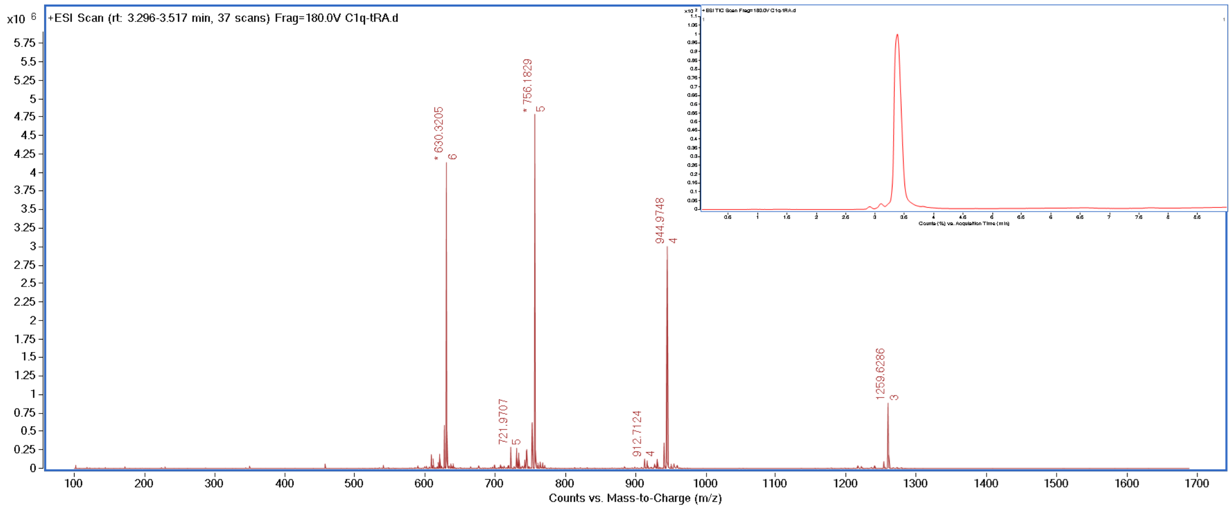


**Supplementary Figure S8.** LC-MS of C1qA-tRA peptide.

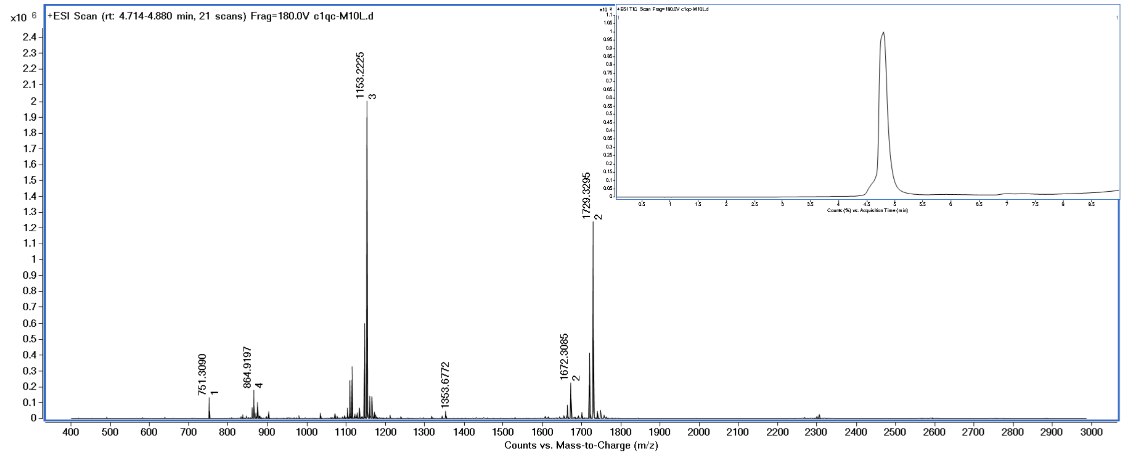


**Supplementary Figure S9.** LC-MS characterization of C1qC-M10L peptide.

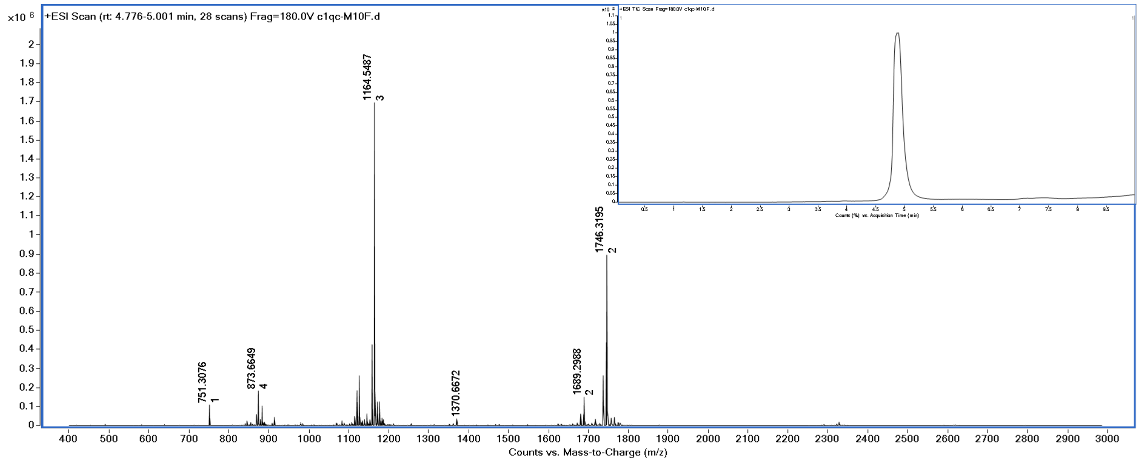


**Supplementary Figure S10.** LC-MS characterization of C1qC-M10F peptide.

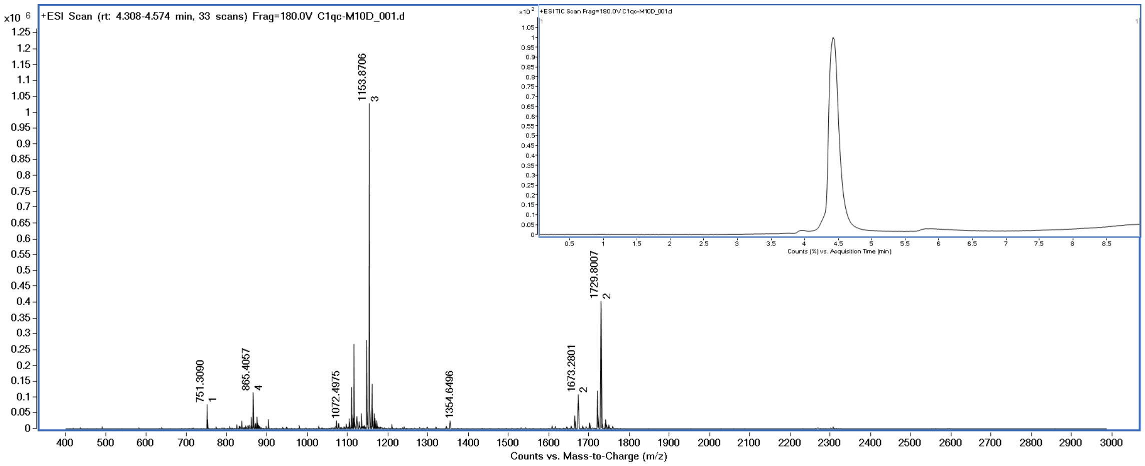


**Supplementary Figure S11.** LC-MS characterization of C1qC-M10D peptide.

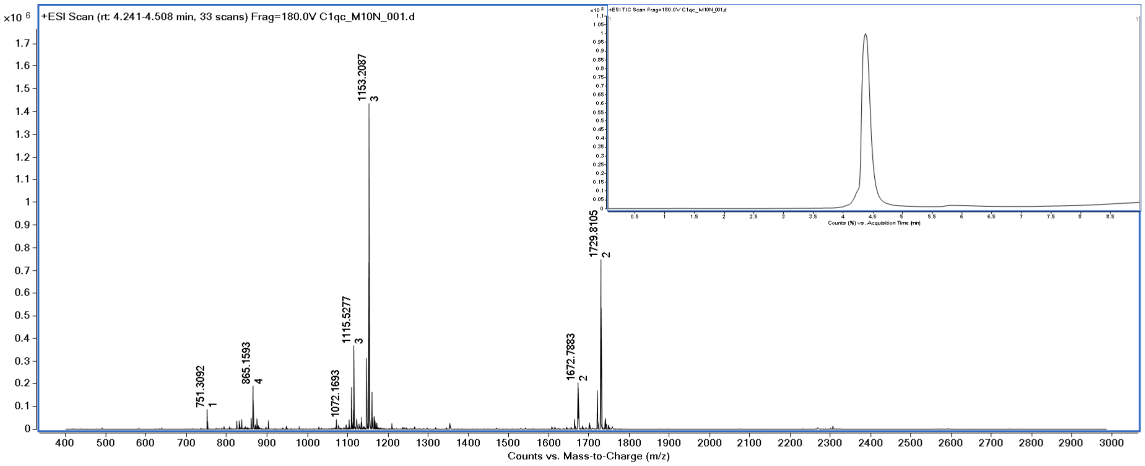


**Supplementary Figure S12.** LC-MS characterization of C1qC-M10N peptide.

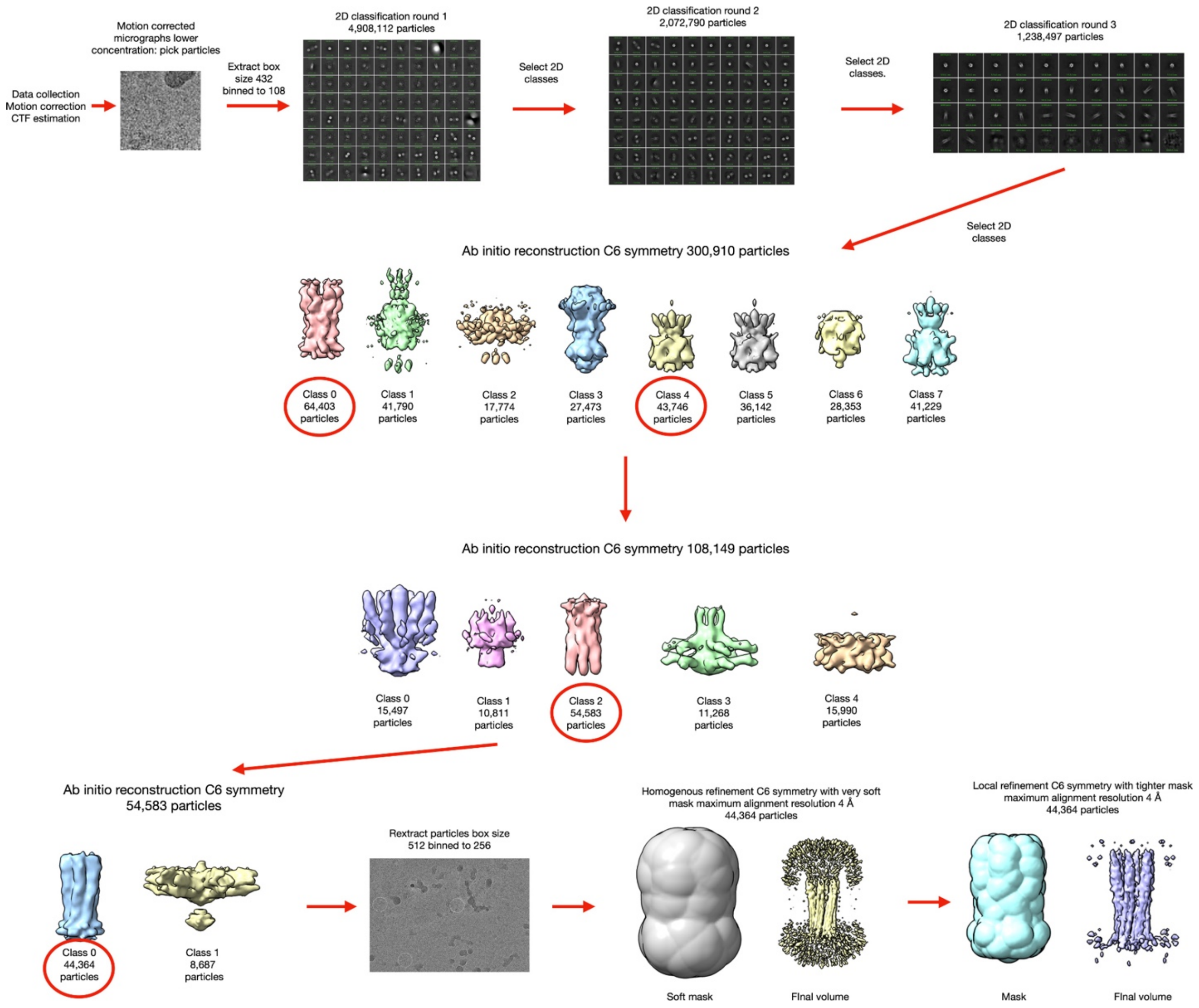


**Supplementary Figure S13.** Workflow for cryo-EM reconstruction of the larger C1q cys crt density map.

**Table S3.** Cryo-EM map and model statistics

| | C1q cys crt Narrow region map + model | C1q cys crt EMReady narrow region map +model | C1q cys crt low resolution full map +model | C1q cys crt low resolution EMReady full map + model | C-M10N just full density map | C-M10F just full density map | hC1q assembly |
|---|---|---|---|---|---|---|---|
| PDB ID | 9C9L | 9C9L | 9C9U | 9C9U | N/A | N/A | N/A |
| EMDB ID | EMD-45363 Additional map | EMD-45363 Primary map | EMD-45371 Primary map | EMD-45371 Additional map | EMD-45373 | EMD-45365 | EMD-45372 |
| map:map 0.143 FSC | 3.7 | N/A | 3.8 | N/A | 4.5 | 4.5 | 3.6# |
| Map:model 0.5 FSC | 3.9 | 3.5 | 5.4 | 5.2 | N/A | N/A | N/A |
| Clash score | 9.15 | 9.15 | 19.72 | 19.72 | N/A | N/A | N/A |
| CC (mask) | 0.77 | 0.83 | 0.78 | 0.62 | N/A | N/A | N/A |
| MolProbity Score | 1.49 | 1.49 | 2.36 | 2.36 | N/A | N/A | N/A |
| Ramachandran favored (%) | 100 | 100 | 89.29 | 89.29 | N/A | N/A | N/A |
| Ramachandran allowed (%) | 0 | 0 | 10.71 | 10.71 | N/A | N/A | N/A |
| Ramachandran outliers (%) | 0 | 0 | 0 | 0 | N/A | N/A | N/A |
| Bond length RMSD (Å) | 0.007 | 0.007 | 0.004 | 0.004 | N/A | N/A | N/A |
| Bond Angle RMSD (°) | 1.258 | 1.258 | 0.987 | 0.987 | N/A | N/A | N/A |
| Rotamer outliers (%) | 0 | 0 | 0 | 0 | N/A | N/A | N/A |
| # particles | 374,558 | 374,558 | 44,364 | 44,364 | 138,738 | 92,001 | 79,400 |
| pixel size | 1.34 (0.67 2x bin) | 1 (EMReady) | 1.34 (0.67 2x bin) | 1 (EMReady) | 1.34 (0.67 2x bin) | 1.34 (0.67 2x bin) | 1.34 (0.67 2x bin) |

*The map model FSC crossed 0.5 FSC twice once at 8.1 Å and again at 4.7 Å. Given the quality of the map and the fact that we could see separated PPII chains (~5 Å apart) we took the 4.7 Å estimate.

#The calculated 0.143 Map:Map FSC is likely an overestimate. We think that the resolution of the map is closer to ~4.5 Å.

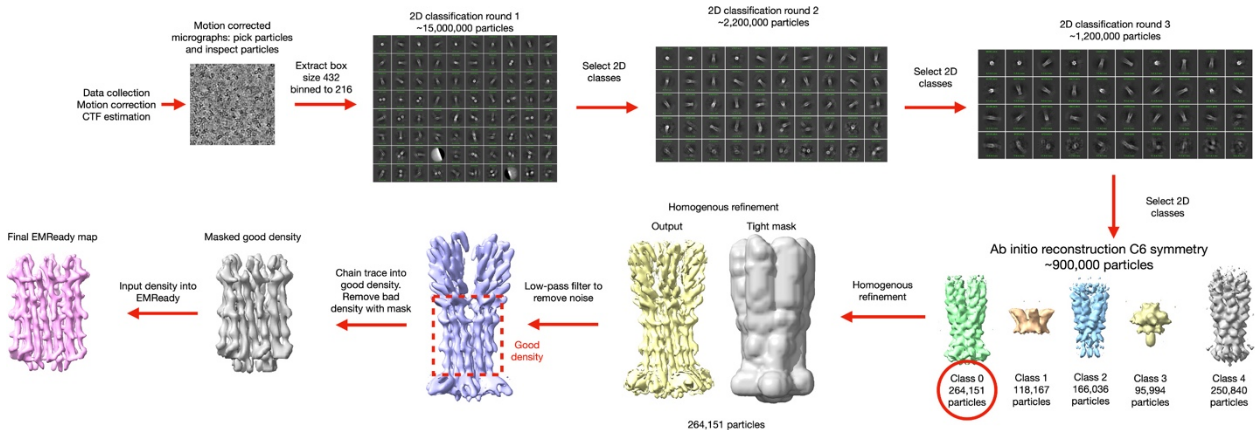


**Supplementary Figure S14.** The general workflow for the processing of cryo-EM data in this paper is shown. In general, all structures determined were processed in this way with differences described in methods.

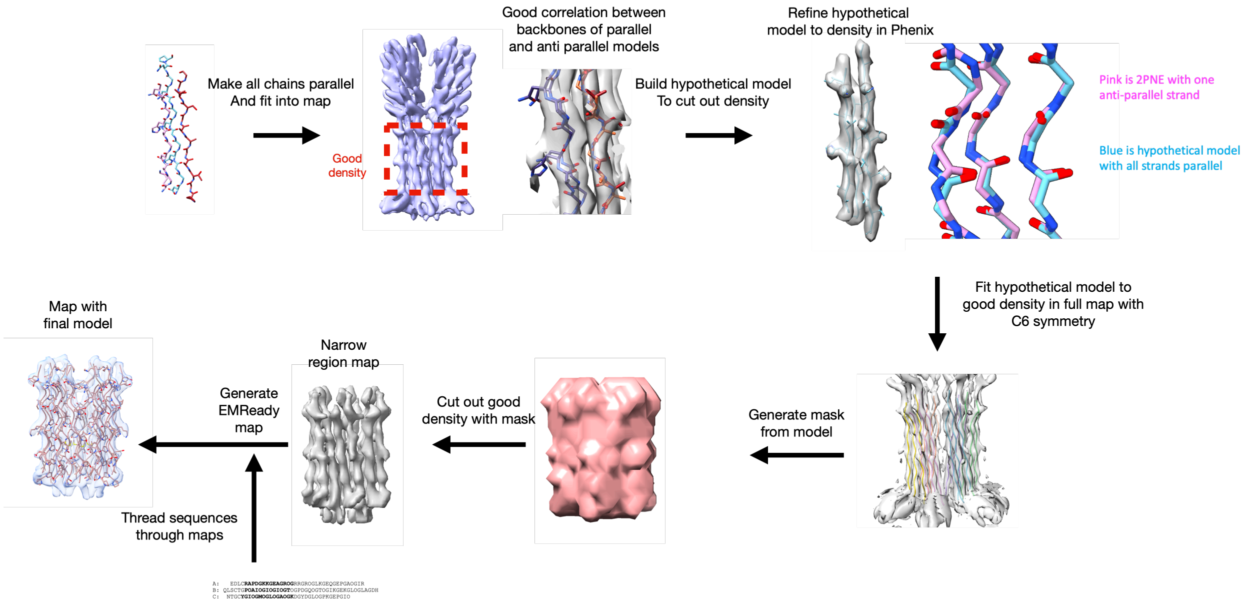


**Supplementary Figure S15.** Description of atomic modeling of narrow region. A general workflow is shown for how the model of the C1q cys crt narrow region was built.

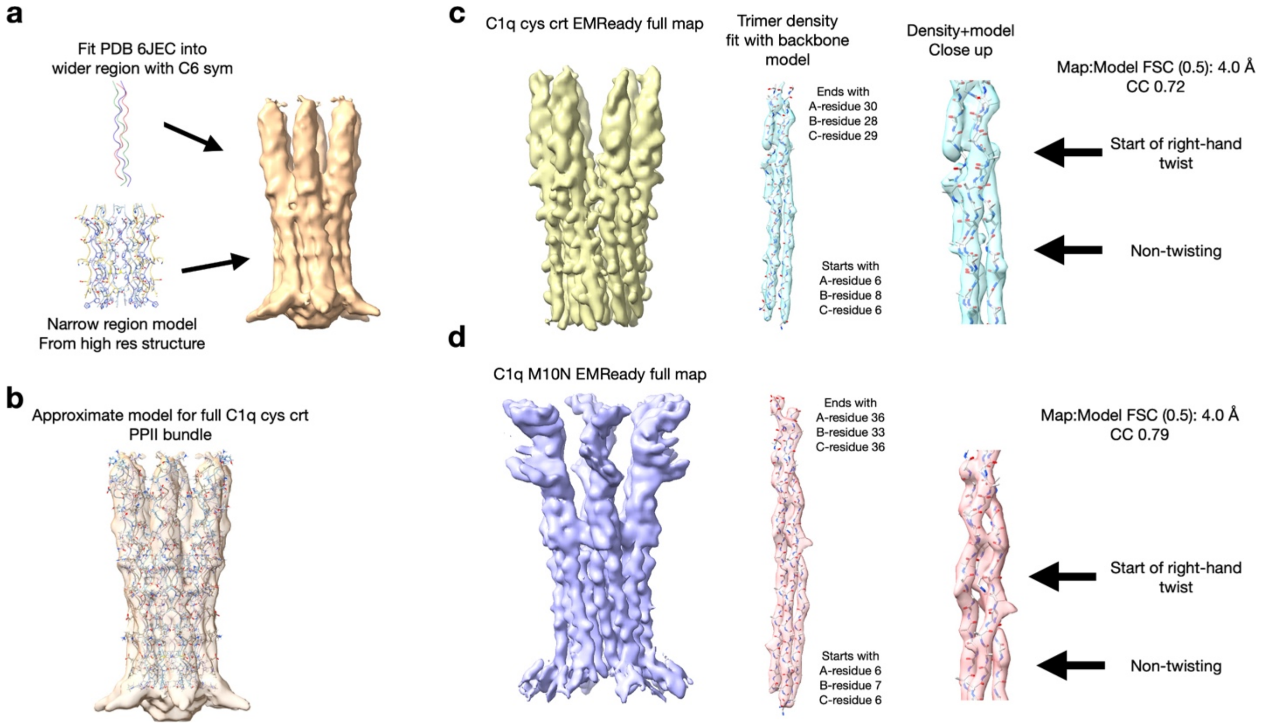


**Supplementary Figure S16.** Description of atomic modeling of the full C1q assembly and fit and comparisons into the EMReady density maps. **a)** Depiction of how the narrow region model and PDB 6JEC were fit into the low-resolution density map. **b)** Fit of the longer C1q cys crt into the low-resolution density map. **c)** EMReady map of the full C1q cys crt structure is shown of the left. In the middle and on the right is a single PPII trimer model fit into its corresponding map. **d**) The EMReady map of the full C-M10N structure is shown on the left. In the middle and on the right is a single PPII trimer model fit into its corresponding C-M10N map.

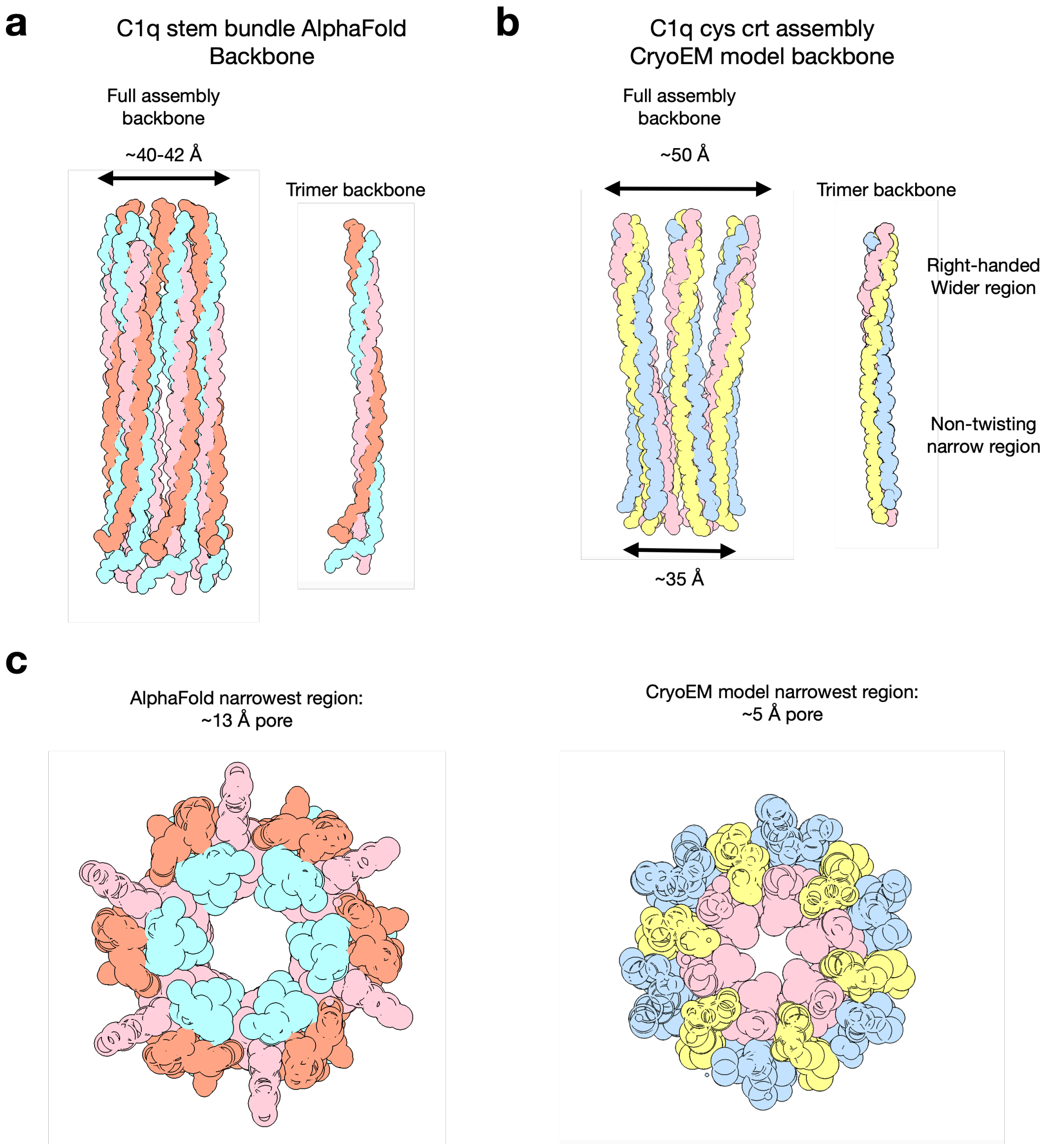


**Supplementary Figure S17.** Comparison between an AlphaFold3 predicted model of the C1q assembly and the C1q cys crt model from cryoEM data. **a)** The left image shows the atomic backbone for the AlphaFold predicted C1q assembly. The right image shows the backbone of a single triple-helical trimer of the assembly predicted by alphafold. **b)** The left image shows the model backbone for the C1q cys crt peptide assembly from the lower resolution full cryo-EM structure. The right image shows a trimer backbone with both the non-twisting PPII helix region as well as the right-handed twisting PPII region. **c)** The left image is a view through the pore of the AlphaFold predicted assembly model showing the ~13 Å wide pore. The right image shows the same view of the C1q cys crt assembly model determined from the cryo-EM data.

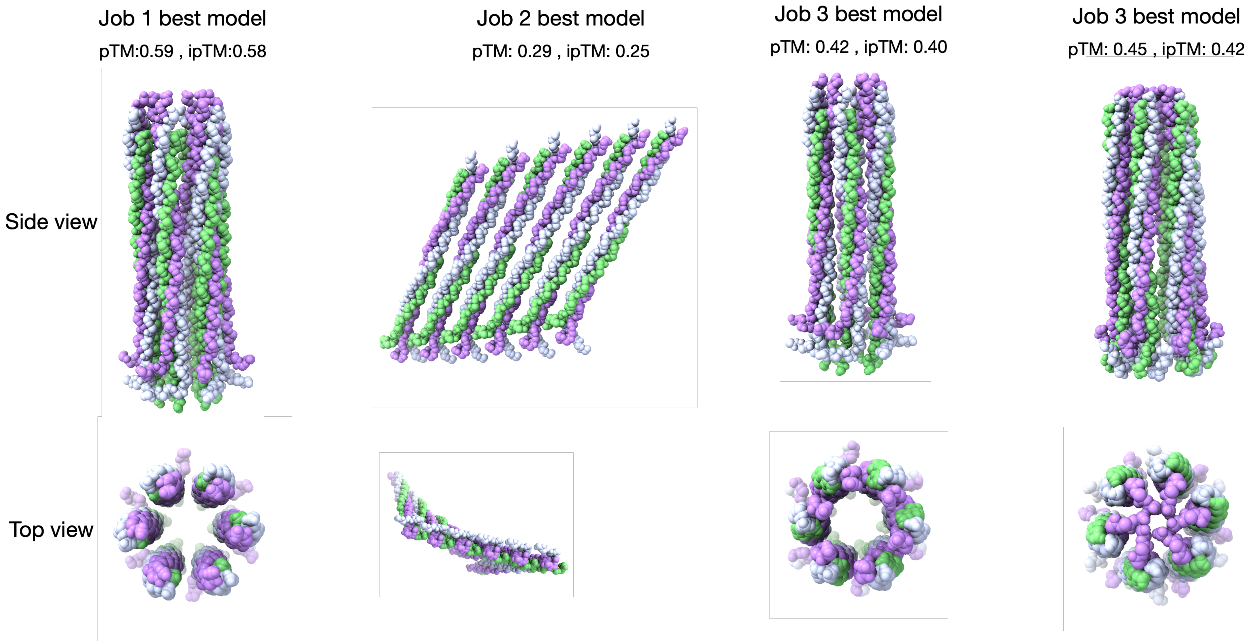


**Supplementary Figure S18.** Comparison of the top scoring models generated from four independent AlphaFold3 jobs on the AlphaFold Server. The term pTM stands for the predicted template modeling score. The term ipTM stands for the interface template modeling score. These are metric used to measure the quality of the overall fold of the complex and the subunit positions in the complex respectively. For pTM greater than 0.5 the overall fold might be similar to the true structure. For ipTM less than 0.6 the prediction is likely failed or incorrect.

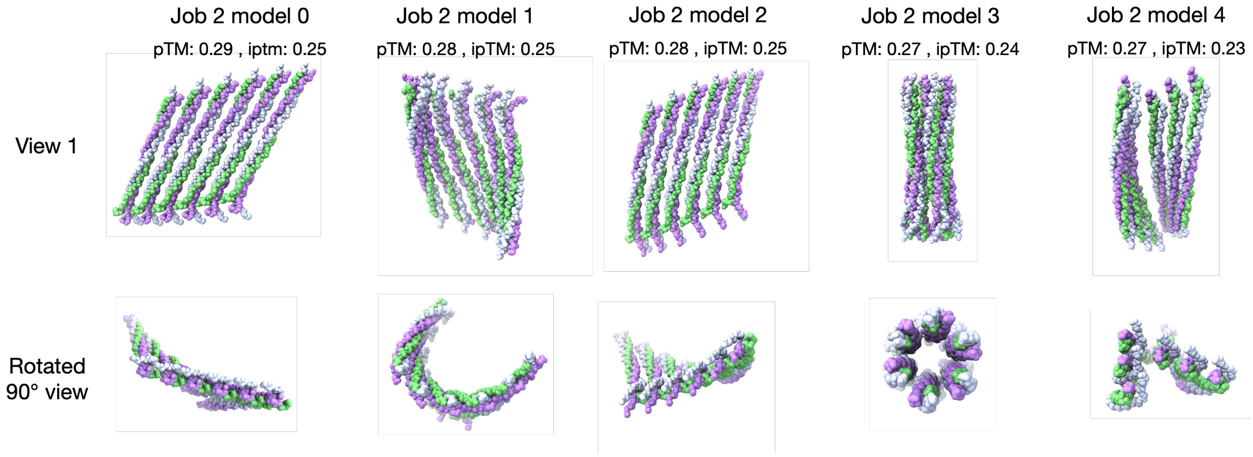


**Supplementary Figure S19.** Comparison of the five models generated from AlphaFold modeling job 2.

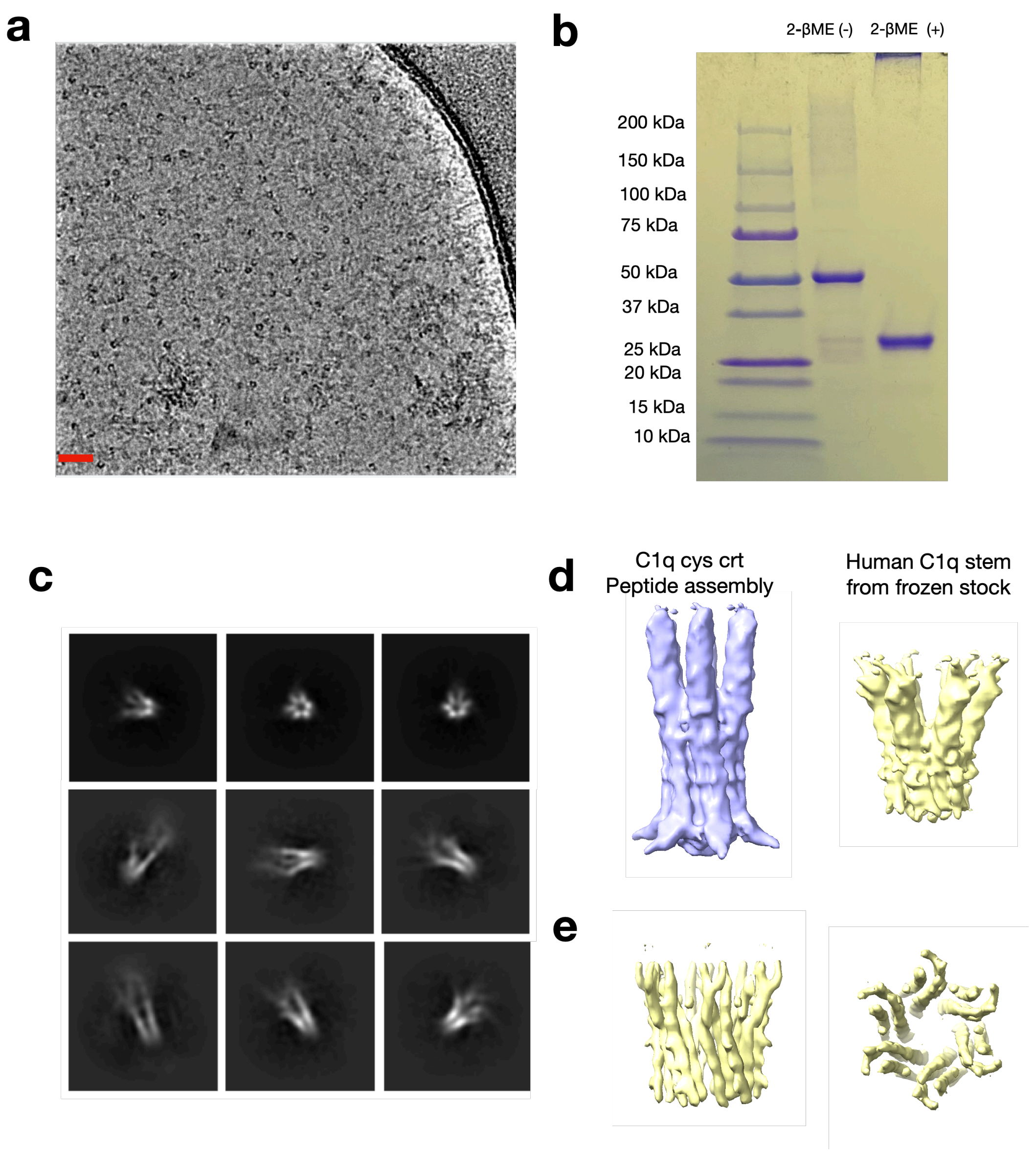


**Supplementary Figure S20.** Cryo-EM structural analysis of the human C1q collagenous stem bundle. a) Cryo-electron micrograph of the hC1q sample. b) SDS-PAGE analysis of the hC1q assembly in the presence, "2-BME(+)", and absence, "2-BME(-)", of reducing agent 2-β-mercaptoethanol. c) 2D-classes of the hC1q stem bundles. e) Comparison in the density maps of the full C1q cys crt peptide assembly (left) and the hC1q stem bundle (right). e) Side view (left) and top view (right) of the resolvable PPII density for the hC1q stem bundle. f) Close-up of the PPII trimer for the hC1q stem bundle showing both the non-twisting region and the region with a right-handed twist.

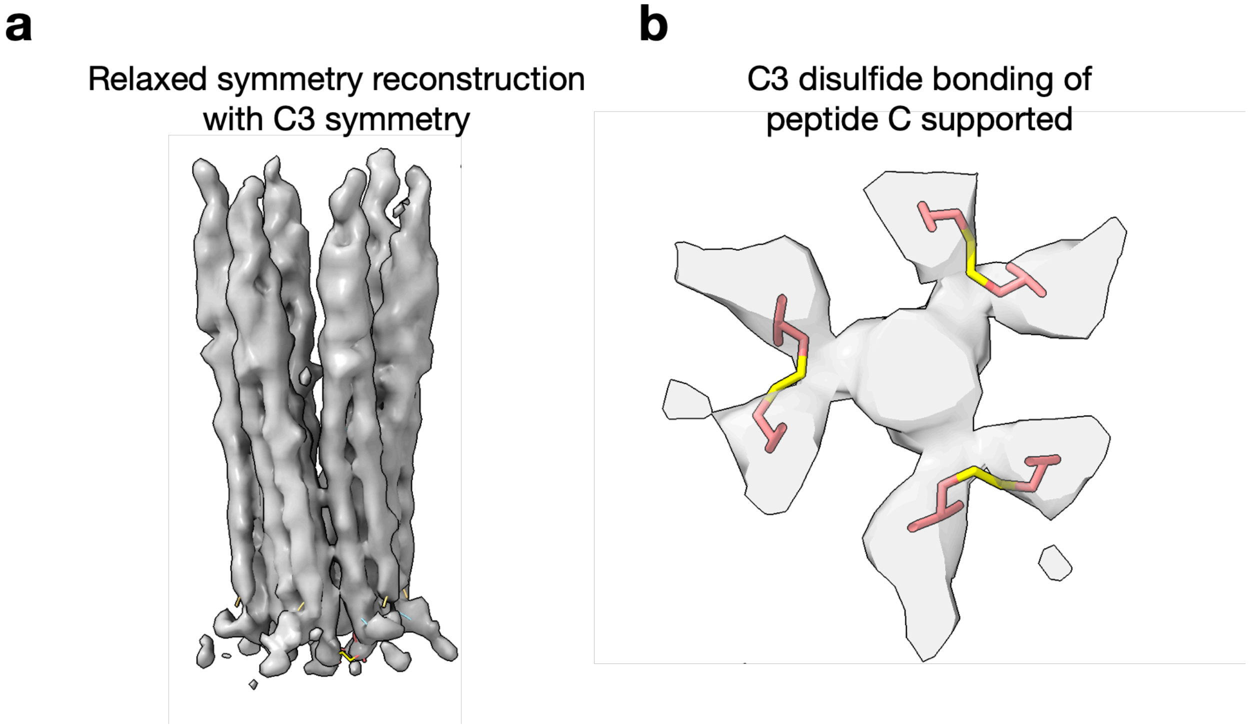


**Supplementary Figure S21.** Likely disulfide bonding scheme in the C1q stem. a) The volume from a relaxed symmetry reconstruction where the C6 symmetry reconstruction and particles were subject to local refinement imposing C3 symmetry. b) Density region of the C3 reconstruction map that appears to be in a plausible location for peptide C disulfide bonding.

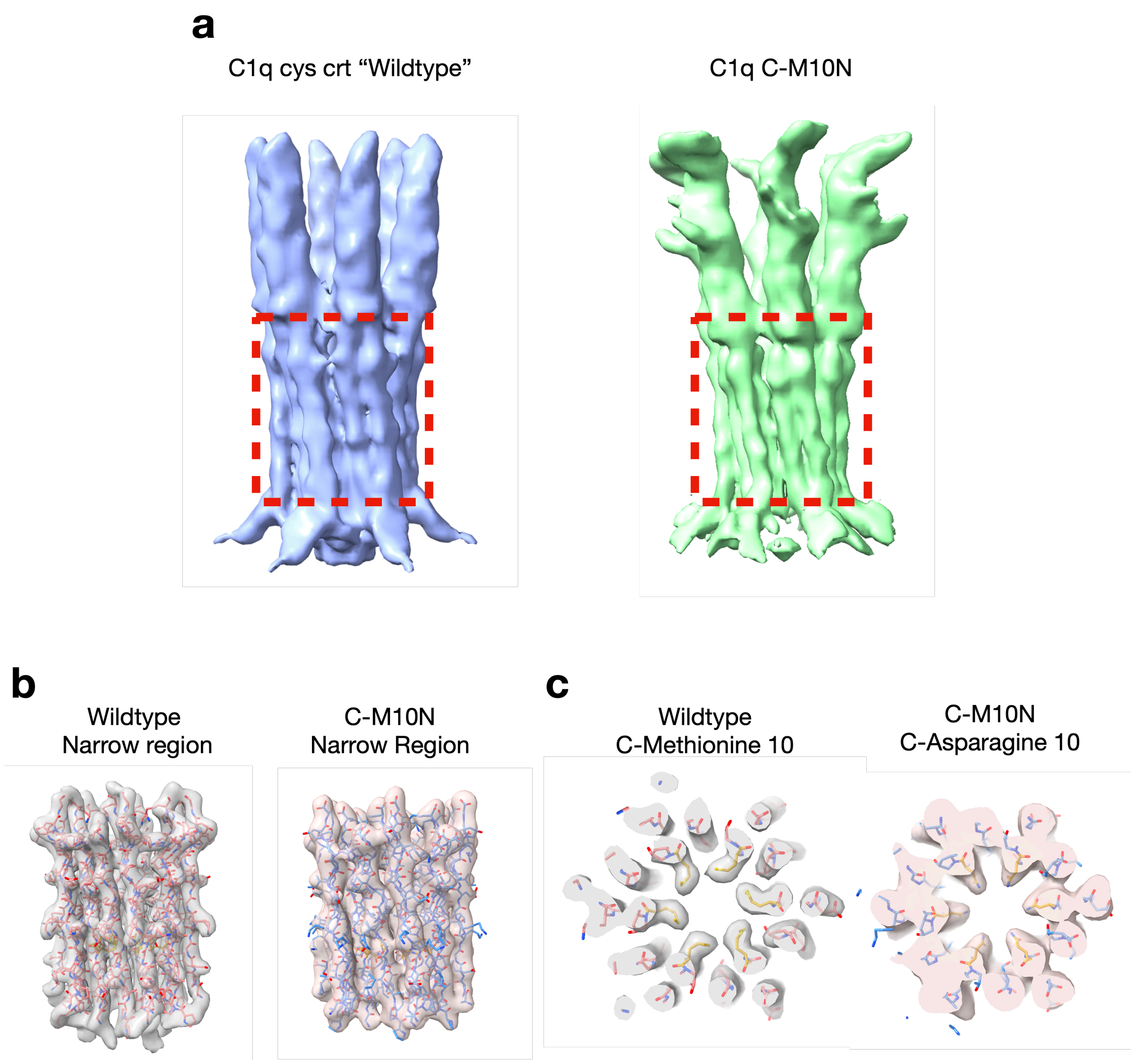


**Supplementary Figure S22.** Comparison of the C1q cys crt (wildtype) and C1q C-M10 cryo-EM structures. a) On the left the low resolution full C1q cys crt density map is shown. On the right the C1q C-M10N density map is shown. b) Comparison of the side views of the cryoEM structures and models of the narrow region for the wildtype (left) and C-M10N (right) assemblies. c) Top-view comparison of the cryoEM density maps and atomic models for wildtype (left) and C-M10N (right) assemblies in the region corresponding to chain C residue 10.

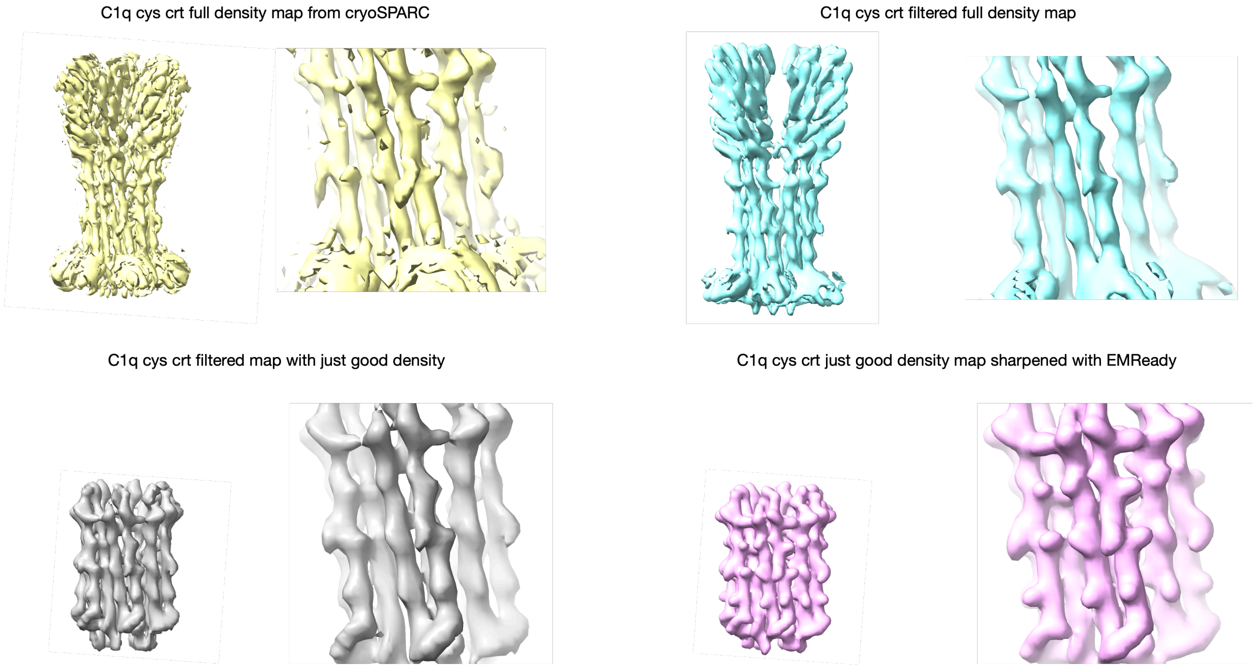


**Supplementary Figure S23.** Comparison of C1q cys crt density maps after cryoSPARC (top-left), after low-pass filtering the cryoSPARC volume (top-right), after cutting out the good density from the narrow region (bottom-left), and after sharpening the narrow region density map with EMReady (bottom-right). For each map the left image shows the full density map while the right image shows a close-up on side chain features.

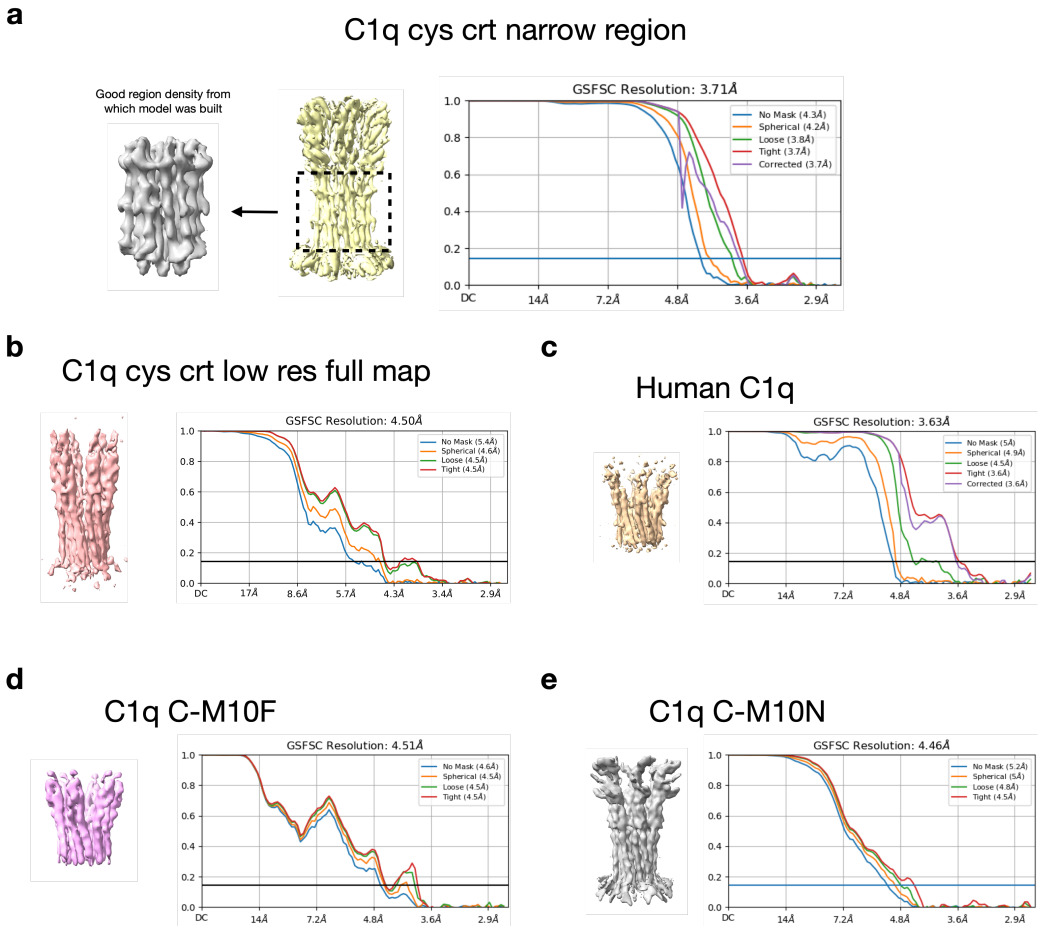


**Supplementary Figure S24.** Map:Map Fourier shell correlation (FSC) curves for each cryo-EM structure solved in this study. a) Map:Map FSC curve for the reconstruction used to generate the C1q cys crt narrow region map and model. The full density map is shown in yellow feature regions of good density outlined by the black dashed line and shown further in the grey density map. **b)** Map:Map FSC curve for the lower resolution C1q cys crt full density map. **c)** Map:Map FSC curve for the human C1q cryo-EM structure. Note that the 3.6 Å Gold-Standard 0.143 FSC threshold appears to be an over estimate of the resolution. For deposition purposes with this map, we used the 0.5 FSC threshold resolution estimate of 4.5 Å. **d)** Map:Map FSC curve for the C1q C-M10F structure. **e)** Map:Map FSC curves for the C1q C-M10N structure.

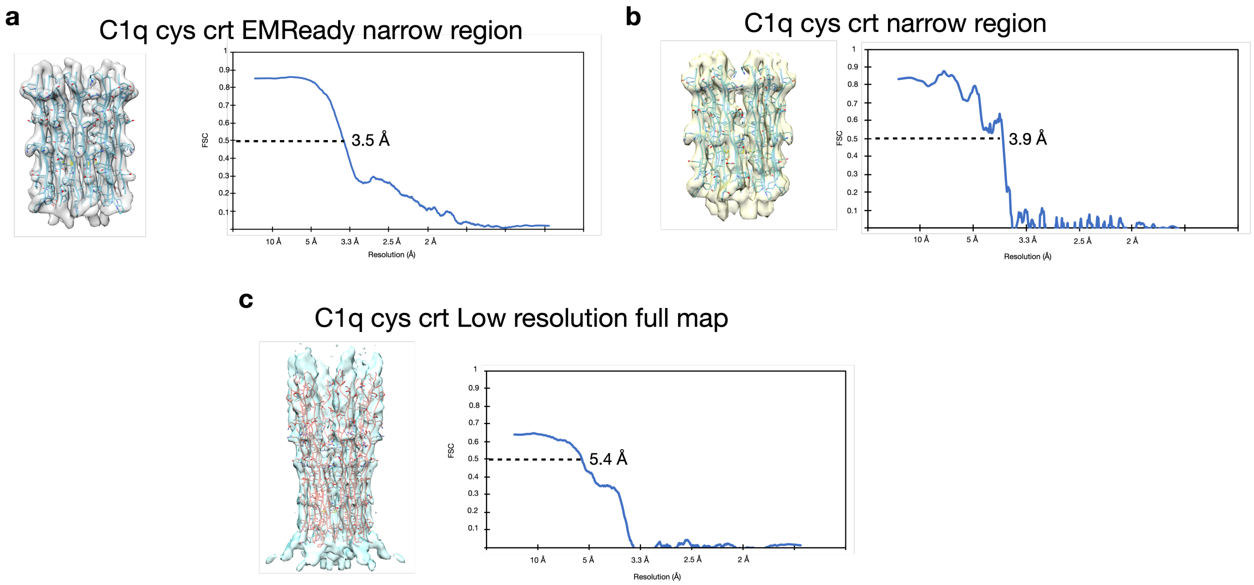


**Supplementary Figure S25.** Map:Model FSC resolution estimates of the structures with models in this study. The map:model FSC curves are shown for a) the EMReady sharpened density map of the C1q cys crt narrow region, b) the C1q cys crt narrow region map prior to EMReady sharpening, and c) the low resolution C1q cys crt full density map and the extended model.

**Additional Files Provided:**

**Supplementary Movie S1.** "C1q_Movie_S1.avi" Provides side views of cryoSPARC 3D variability analysis output.

**Supplementary Movie S2.** "C1q_Movie_S2.avi" Provides top views of cryoSPARC 3D variability analysis output.
